# The nanoscale mobility of calcium channels is driven by readily releasable synaptic vesicles to support precise neurotransmission in live *C. elegans*

**DOI:** 10.64898/2026.03.29.715107

**Authors:** Yunke Zhao, Dai Zhai, Fabien Pinaud

## Abstract

Precise neurotransmission requires voltage-gated calcium channels (VGCCs) to be tightly coupled with synaptic vesicles (SVs) at active zones (AZs). As VGCCs are mobile in the fluid presynaptic membrane, it remained unclear how they spatiotemporally align with SVs for reliable synaptic release. Using super-resolution imaging and single-molecule tracking in *C. elegans*, we report that two diffusive behaviors of N-type VGCCs at AZs are governed by primed SVs via distinct mechanisms. VGCCs diffuse slowly in multiple nanodomains (∼100 nm) that spatially correlate with UNC-10/RIM, and the channels’ mobility and nanodomain numbers are coupled to SNAREs assembly and SV priming levels via RAB-3/UNC-10/UNC-13L tripartite complexes. Outside nanodomains, a faster mobility of VGCCs and their AZ distribution scale with UNC-13S-mediated priming under an antagonistic regulation by UNC-10 and TOM-1/Tomosyn. These findings reveal unexpected mechanisms whereby pools of readily releasable SVs actively control the nanoscale mobility and nanodomain organization of VGCCs to support precise neurotransmission.

## Introduction

At the presynaptic active zone (AZ), exocytosis of synaptic vesicles (SVs) is spatiotemporally orchestrated by an evolutionarily conserved release machinery^1–3^ that includes multi-subunits voltage-gated calcium channels (VGCCs), which provide crucial calcium influx for SV release^4,5^. In both mammals and *C. elegans* nematodes, VGCCs are recruited in close proximity to SVs mainly by the AZ modulator UNC-10/RIM, which interacts with the channel’s α1 pore-forming subunit UNC-2/Ca_v_2 and binds vesicular Rab-3 to dock SVs^6–9^. UNC-10/RIM also directly regulates UNC-13/Munc-13, which primes SVs to become readily releasable^10–12^. During priming, UNC-13/Munc-13 tethers SVs to the plasma membrane and coordinates the assembly of SNARE complexes by conformationally opening and aligning UNC-64/Syntaxin-1 (t-SNARE) with both SNAP-25 (t-SNARE) and vesicular SNB-1/VAMP (v-SNARE), which drive SV fusion upon calcium entry^13–16^. By interacting with UNC-13/Munc-13, UNC-10/RIM couples VGCCs to readily releasable SVs, awaiting calcium influx^17^. The local density of VGCCs and their coupling distances to SVs are therefore key determinants of the SV release probability and its efficacy^18,19^. Indeed, synapses having slow release kinetics are characterized by a relatively broad, microdomain coupling between VGCCs and SVs (>100 nm), while fast-releasing synapses adopt a much narrower, nanodomain coupling (<100 nm)^20,21^.

While precise spatial coupling between VGCCs and SVs is critical for efficient neurotransmissions, maintaining such nanoscale proximities is complicated by the lateral diffusion of VGCCs in the fluid presynaptic membrane, as observed in diverse synaptic systems^22–24^. Indeed, the mobility of VGCCs at AZs has emerged as a new determinant of presynaptic release, with VGCC mobilities scaling with intracellular calcium levels and neural activities, and alteration of channel diffusion shaping short-term synaptic plasticity and homeostasis^23–25^. Because VGCCs are not static, one would expect their coupling distances to SVs and release probabilities to shift continuously. How the AZ organization maintains precise spatiotemporal coupling between mobile VGCCs and SVs for reliable synaptic releases remains unclear.

Here, we describe how readily releasable SVs actively drive the nanoscale mobility and spatial organization of N-type VGCCs within intact neuromuscular synapses of live *C. elegans*. Using single particle tracking and super-resolution imaging, we characterized two distinct diffusive behaviors of VGCCs at AZ. We show that VGCCs diffuse slowly within multiple intra-AZ nanodomains (∼100 nm) that are organized in the typical dense-projection-like landscape of *C. elegans* and that spatially correlate with UNC-10. Outside these nanodomains, the channels adopt a faster diffusion mode while being confined within the larger, diffraction-limited AZ puncta (∼300 nm). We further show that the two mobilities of VGCCs are governed by SV priming and SNARE complex assembly under the coordination of UNC-10, but by different UNC-13 isoforms. Specifically, the slow mobility and nanodomain number of VGCCs are predominantly regulated by UNC-13L-mediated priming through tripartite SV coupling complexes involving RAB-3, UNC-10 and UNC-13L, while the fast mobility and AZ distribution of VGCCs outside nanodomains scale with the pool of UNC-13S-primed SVs under antagonistic regulations by UNC-10 and the t-SNAREs inhibitor TOM-1/Tomosyn. We propose that the heterogeneous mobilities and spatial organizations of N-type VGCCs provide distinct channel-vesicle coupling strategies for different pools of readily releasable SVs, offering a tunable mechanism to set SV release probabilities.

## Result

### N-type VGCCs exhibit two distinct confined diffusive behaviors in live *C. elegans*

At neuromuscular junction (NMJ) synapses in *C. elegans*, the single N- and P/Q-type VGCCs (UNC-2/Cav2) mediate both evoked and tonic synaptic releases^26^. The functions and surface trafficking of UNC-2/Cav2 require the extracellular α2δ subunit UNC-36, whose own AZ localization also depends on UNC-2/Cav2^26–28^. To study the dynamic distributions and nanoscale mobilities of VGCC complexes, we used complementation activated light microscopy (CALM) with single molecule sensitivity in intact living *C. elegans*^29,30^. We endogenously tagged UNC-36/α2δ with a N-terminal dark split sfCherry1-10^31^ (sCherry-UNC-36) using CRISPR/Cas9^32^. To activate the fluorescent signals of the extracellular-facing sCherry-UNC-36, we introduced a synthetic complementary peptide (sfCherry11) via pseudocoelomic microinjections^29,30^, which allows for specific detections of surface VGCCs without interference from intracellular background signals (Fig. 1a). We previously reported that N-terminal tagging of split-fluorescent proteins to UNC-36/α2δ does not alter the conducting activities of VGCCs^29^ and animals expressing sCherry-UNC-36 had functional channels and exhibited normal locomotion compared to N2 Bristol wild-type (WT) animals^33^ (Fig. S1a). After fluorescence activation by CALM, surface sCherry-UNC-36 showed puncta localization along the nerve cords as reported^29^ and colocalized with endogenous UNC-2/Cav2^27^ (Fig. 1b; Fig S1b). To further confirm the AZ localization of sCherry-UNC-36, we endogenously tagged the AZ marker UNC-10/RIM with 5xMyc and UNC-2/Cav2 with HA at the N-terminus. Those transgenic animals showed normal locomotion and synaptic release compared to WT (Fig. S1c and S1d). Consistent with previous findings^9,27,28^, N-type VGCCs puncta represented by either surface sCherry-UNC-36 or HA-UNC-2 colocalized with Myc-UNC-10 puncta at AZs, and overlapped with NMJ boutons along the nerve cords (Fig. S1e and S1f).

**Figure 1.**
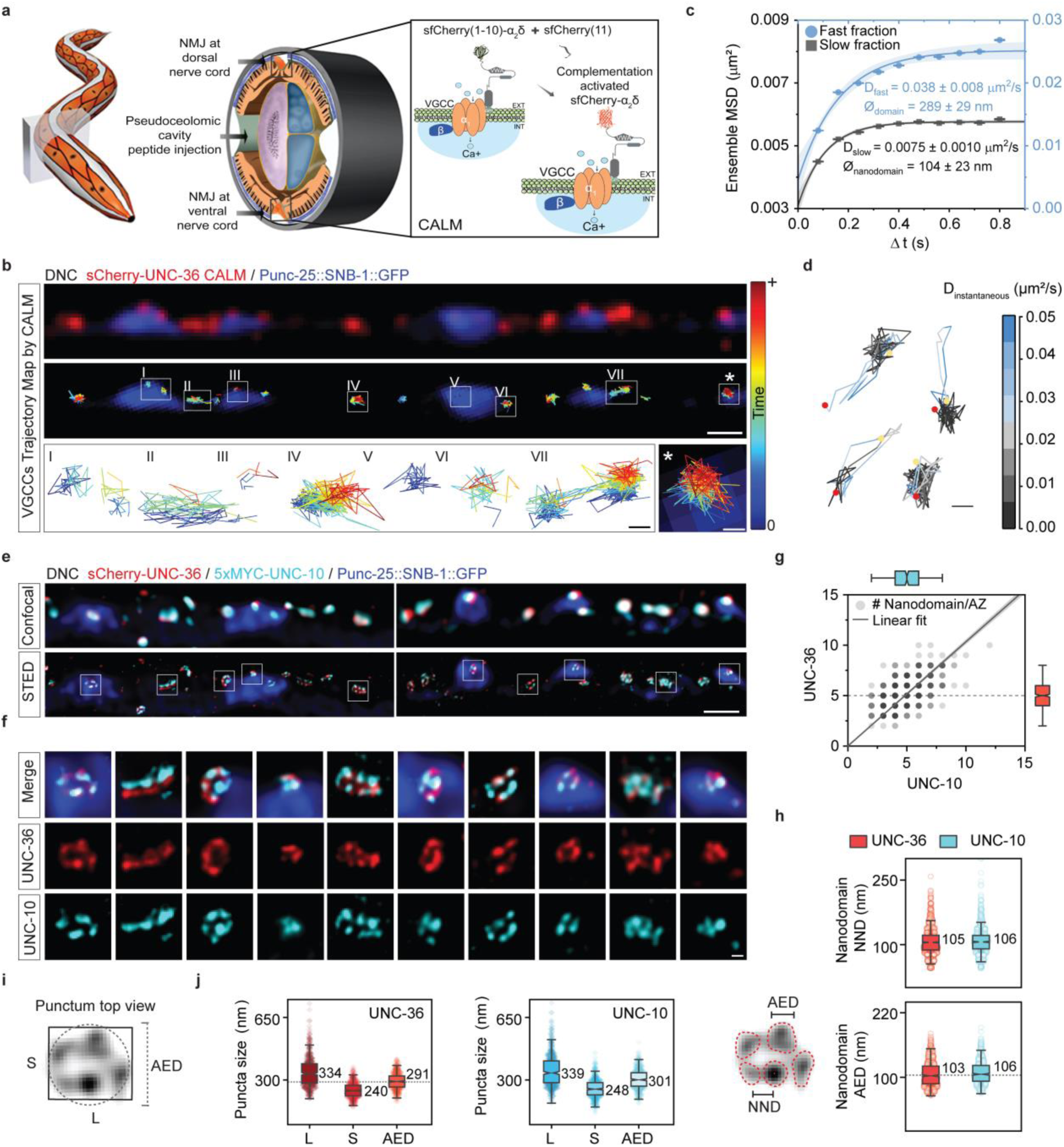
N-type VGCCs exhibit two distinct diffusive behaviors within multiple nanodomains that correlate with UNC-10 at AZ. **(a)** Schematic of CALM by pseudocoelomic microinjection of synthetic sfcherry (11) peptides to activate extracellular split sfCherry (1-10) tagged to UNC-36 and detect surface-localized VGCC complexes at *C. elegans* NMJ. Adapted with permission from ref.^42^ and from Z. F. Altun and D. H. Hall (wormatlas.org)^43^. **(b)** Sum projection of a dual-color SM CALM movie by HILO light sheet imaging of sCherry-UNC-36 (red) along a dorsal nerve cord (DNC) marked with GABAergic Punc-25::SNB-1::GFP (blue, top). Corresponding VGCC trajectory maps color-coded by time (Tmin = blue; Tmax = red, middle) and zoom of selected insets (bottom). Scale bars, 1 µm (middle) and 100 nm (bottom). **(c)** Ensemble mean square displacement (MSD ± SEM) of slow (black squares) and fast (blue dots) fractions for VGCCs (n = 42,391; 58 animals). MSD curves are fitted with a confined 2D diffusion model and the fit 95% confidence bands are shown together with the diffusion coefficient (D ± SEM) and the domain diameter (∅ ± localization precision) for each fraction. **(d)** Representative trajectories of heterogeneous VGCC diffusive behaviors color-coded by instantaneous diffusion coefficients (D_instantaneous_). Trajectory start: yellow dot; trajectory stop: red dot; scale bar: 100 nm. **(e)** Confocal and dual-color STED images of immunostained surface sCherry-UNC-36 (red) and 5xMYC-UNC-10 (cyan) along dorsal nerve cords (DNC) marked by Punc-25::SNB-1::GFP (blue). Scale bars, 1 µm. **(f)** Zoomed insets of STED images in **(e)**. Scale bars, 100 nm. **(g)** Correlation between the number of UNC-36 (red) and UNC-10 (cyan) nanodomains per AZ (n = 516; 10 animals). Linear regression indicates proportional scaling (y=1.03x, R^2^: 0.932). Notched boxes: median ± interquartile range (IQR); whiskers: 1.5xIQR. Dashed line marks the median number of UNC-36 nanodomains. **(h)** Nearest-neighbor distance (NND) and area-equivalent circular diameter (AED) of nanodomains for UNC-36 (red; n =1416 NND; 993 AED, 10 animals) and UNC-10 (cyan; n = 1454 NND; 786 AED; 10 animals). Notched boxes: median ± IQR; whiskers: 1.5xIQR. Dashed line marks the confinement size of the slow regime for VGCCs in (C). **(i, j)** Puncta sizes of UNC-36 (red) and UNC-10 (cyan) (n = 516; 10 animals). Notched boxes: median ± IQR; whiskers: 1.5xIQR. Dashed line marks the confinement size of the fast regime for VGCCs in (c).

In *C. elegans*, UNC-36/α2δ is also required for the function of L-type Egl-19/Cav1 channels in muscles, but is dispensable for Egl-19/Cav1-mediated tonic releases at NMJs^26,34^. To test whether synaptic sCherry-UNC-36 also represents L-type VGCCs, we probed the extent to which UNC-36/α2δ colocalizes with Egl-19/Cav-1 at nerve cords versus muscles. In contrast to UNC-2/Cav2, endogenous Egl-19/Cav1^35^ showed a broad distribution along nerve cords, with little co-localization with either GFP-UNC-2 or surface sCherry-UNC-36 (Fig. S1f-S1h). This is consistent with previous findings that UNC-2/Cav2 and Egl-19/Cav1 localize to distinct synaptic regions^36,37^ and aligns with the reported function of UNC-36/α2δ, which is required for UNC-2/Cav2 but not Egl-19/Cav1 mediated synaptic releases at NMJs^26^. The dispersed distribution of neuronal Egl-19/Cav1 was also in stark contrast to the puncta distribution of muscular Egl-19/Cav1, which extensively colocalized with surface sCherry-UNC-36 at the sarcolemma of muscles^29,30,34^ (Fig. S1i and S1j). Together, these results indicate that UNC-36/α2δ is predominantly associated with UNC-2/Cav2 at AZs rather than with Egl-19/Cav1.

To track individual VGCCs by CALM, the sub-diffraction localizations of single sCherry-UNC-36 signals were extracted with a precision of ∼20 nm and linked frame to frame to construct the diffusion trajectory^38^ of each channel (Fig. 1b; Fig. S2a). At AZs, individual VGCCs exhibited heterogeneous diffusion dynamics with single-step sizes ranging from a few tens to hundreds of nanometers per frame (Fig. 1b; Fig. S2b; Movie S1-S2). To quantitatively define these different mobilities, we analyzed pooled trajectories (n=42,391 from 58 animals) based on the probability distribution of their squared displacements^29,30,39^. We found that VGCCs have two distinct confined diffusive behaviors at AZs in live *C. elegans* (Fig. 1c). The major fraction of VGCC (81%) exhibited a slow mobility, with a diffusion coefficient D_slow_ of 0.0075 ± 0.0010 μm^2^/s and were confined within circular nanodomains 104 ± 23 nm in diameter. The remaining 19% of VGCCs displayed a five-fold faster diffusion (D_fast_ = 0.038 ± 0.008 μm^2^/s) and were confined within larger domains 289 ± 29 nm in diameter (Fig. 1c). Within individual VGCC trajectories, alternations between slow and fast diffusions were observed (Fig. 1d; Fig. S2c), indicating that D_slow_ and D_fast_ represent two diffusive behaviors of single channels rather than two independent populations of VGCCs. Later observations also show that removal of AZ proteins that directly interact with UNC-2 alters both diffusions, supporting that D_fast_ reflects a genuine faster mobility of VGCC complexes rather than a transient dissociation of UNC-36 from UNC-2. While these alternating diffusive behaviors of VGCCs in *C. elegans* are unique, their respective diffusion coefficients are similar to those of Cav2 in different organisms. For instance, D_slow_ matches the mean diffusion coefficient reported for Cac/Cav2 in *Drosophila* NMJs (0.0075 μm^2^/s)^25^, while D_fast_ is similar to the mobility reported for Cav2.1 in rodent hippocampal synapses (0.04 μm^2^/s)^23^. The small confinement size of VGCCs falls between those measured for rodent Cav2.1 (∼80 nm)^23^ and Drosophila Cac/Cav2 (∼117 nm)^25^, while their larger confinement matches the expected size of AZ in *C. elegans*^40^. This suggests that the nanoscale mobilities of VGCCs reflect intrinsic structural features of AZs^41^, which we studied next.

### N-type VGCCs diffuse in multi-nanodomains that spatially correlate with UNC-10/RIM at AZs

The hallmark of *C. elegans* AZ is an electron dense projection (DP) with branched ultrastructure and ∼ 100 nm bay-like sites for SV docking and release^44,45^. We therefore determined whether the confinement of VGCCs in nanodomains reflects those AZ sub-structures by resolving the nanoscale distribution of the channels and that of UNC-10 using stimulated emission depletion (STED) microscopy. Interestingly, we found that VGCCs and UNC-10 form multiple co-localizing nanodomains within individual AZ puncta, in agreement with our diffusion analyses (Fig. 1e and 1f; Fig. S3a-d). Notably, the spatial organization of these nanodomains closely resembled the bay-like sites between DP branches^44^, consistent with the known functions of UNC-10/RIM in docking SVs and coupling VGCCs^1,3,46^. Similar to the variability in the number of docking bays across different DPs^44^, we also observed heterogeneities in nanodomain numbers per AZ. Most AZs (∼50%) contain 4-5 VGCC or UNC-10 nanodomains, 28-37% contain more than 6 nanodomains, while 15-22% have 1-3 nanodomains (Fig. S3e). Quantitative analyses showed a significant correlation between the numbers of VGCC and UNC-10 nanodomains, both exhibiting a median of five nanodomains per AZ along the dorsal nerve cords (interquartile range [IQR]: 4/6) (Fig.1g). While not strictly identical, the nanodomains of VGCCs and UNC-10 showed similar spatial distribution, with a median nearest neighbor distance (NND) of 105 nm (IQR: 87/122nm) and 106 nm (IQR: 90/121nm), respectively (Fig. 1h). The size of VGCC nanodomains (median Area Equivalent Diameter AED: 103 nm; IQR: 88/121nm) also aligned with the size of UNC-10 nanodomains (106 nm; IQR: 93/122nm) and importantly, closely matched the confinement size of slow-diffusing VGCCs extracted by diffusion analyses (104 nm ± 23 nm) (Fig. 1c and 1h). This indicates that the slow mobility (81%) of VGCCs reflects their diffusion within individual nanodomains that spatially correlate with UNC-10.

Within each AZ, multiple VGCCs or UNC-10 nanodomains form an elliptic punctum similar to the overall DP geometry established by electron microscopy^44,45^ (Fig. 1f). The size of VGCC puncta, defined by their long and short axes (median: 334 x 240 nm), strongly correlates with that of UNC-10 (median: 339 x 248 nm) (Fig.1i and 1j; Fig. S3f). To adequately compare these parameters with the circular confinement size of fast-diffusing VGCCs established by diffusion analyses, we also calculated their area equivalent diameter (AED). The median AED of VGCCs was 291 nm (IQR: 260/322nm) and that of UNC-10 was 301 nm (IQR: 265/340nm) (Fig. 1j). Both sizes are consistent with the confined domain (289 ± 29 nm) of fast-diffusing VGCCs, suggesting that the fast mobility (19%) reflects their exploration across the multi-nanodomain AZ area.

### UNC-10 enhances the diffusion of VGCCs and promotes their multi-nanodomain organization

To investigate whether VGCC nanodomains are organized by UNC-10, we examined their nanoscale distribution in an *unc-10(md1117)*^8^ null mutant. *Unc-10(md1117)* animals showed severely impaired neurotransmission and fewer VGCC puncta along the nerve cords as previously described^9,47^ (Fig. 2a; Fig. S4a-4c). By STED, we found that, in the absence of UNC-10, VGCC nanodomains persist (Fig. 2b), but their median number decreases significantly from 5 to 2 per punctum (Fig. 2c). The percentage of VGCC puncta containing more than 3 nanodomains dropped from 85% in WT animals to 11% in *unc-10(md1117)* mutants (Fig. 2d). Meanwhile, the NND between VGCC nanodomains also significantly increased, resulting in a more dispersed organization compared to WT (Fig. 2e). These results show that, while VGCC nanodomains form independently of UNC-10, UNC-10 promotes their accumulation at AZs and maintains them in a tight spatial organization.

**Figure 2.**
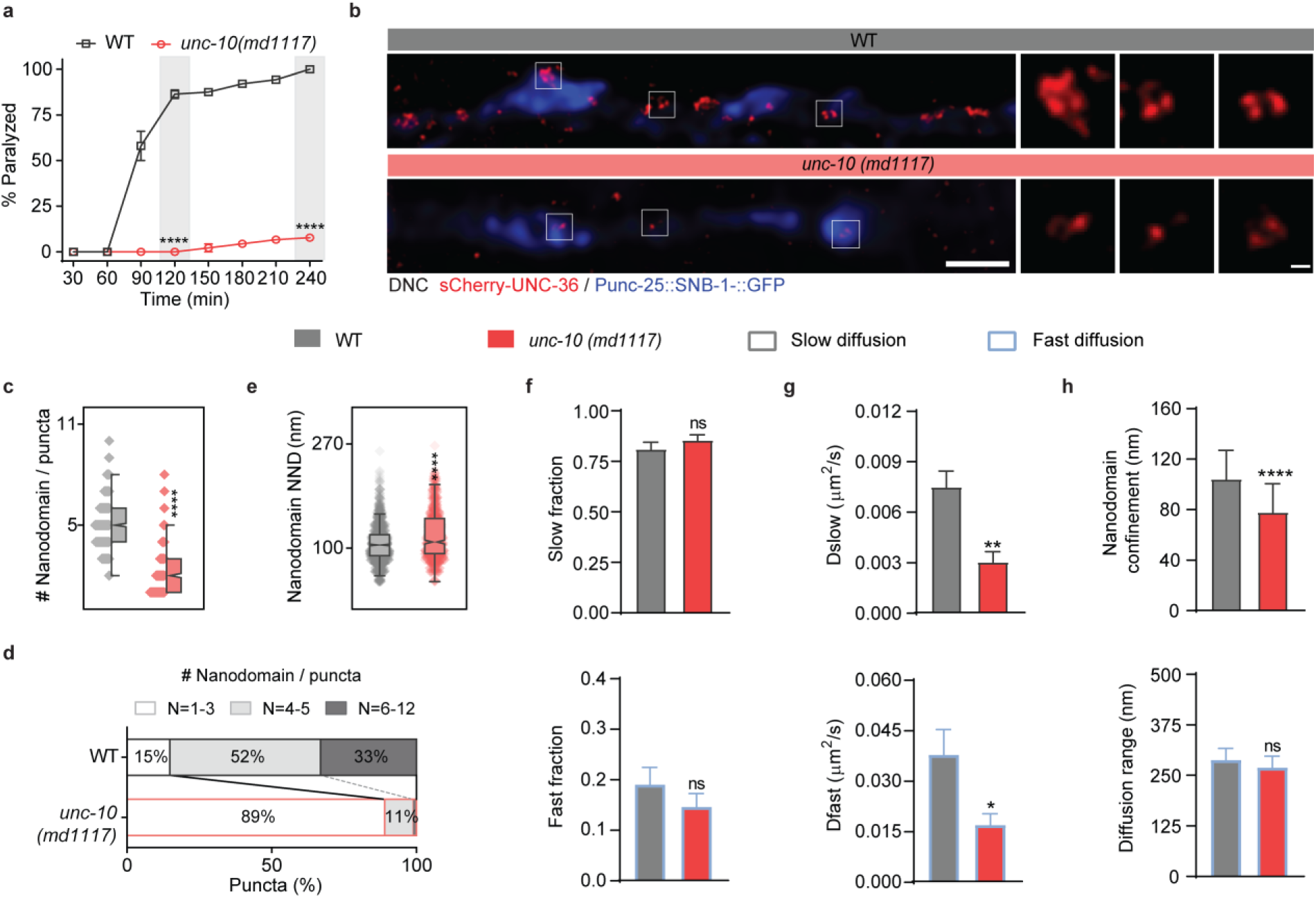
UNC-10 enhances VGCCs diffusion and promotes the multi-nanodomain organization of VGCC. **(a)** Percentage of paralyzed animals over time (mean ± SEM) in aldicarb-sensitivity assays, comparing WT and *unc-10(md1117)* genotypes (n = 90). Two-way ANOVA and Tukey’s tests, ****p<0.0001. **(b)** Representative STED images of immunostained surface sCherry-UNC-36 (red) overlayed with confocal images of Punc-25::SNB-1::GFP (blue) at the dorsal nerve cords of WT or *unc-10(md1117)* animals. Scale bars, 1 µm and 100 nm for insets. **(c-e)** Number (c), distribution (d) and nearest-neighbor distance NND (e) of UNC-36 nanodomains per diffraction-limited puncta in WT (gray; n = 1076 puncta; 1416 NND; 10 animals) and *unc-10 (md1117)* mutants (red; n = 634 puncta; 855 NND; 7 animals). Notched boxes: median ± interquartile range (IQR); whiskers: 1.5xIQR. Comparisons to WT by Kruskal-Wallis and Dunn’s test, ****p < 0.0001. **(f-h)** Mobility of VGCCs in WT (gray; n= 42,391 trajectories; 58 animals) and *unc-10 (md1117)* mutant (red; n= 18,645 trajectories; 34 animals) with fractions (f, mean ± SEM), diffusion coefficients (g, mean ± SEM), and nanodomain confinement size or diffusion range (h, mean diameter ± localization precision) for the channels’ slow (top row, black) and fast (bottom row, blue) diffusive behaviors. Comparisons to WT by two-tailed T-test. ****p < 0.0001; **p<0.01; *p<0.05; ns, not significant.

We then studied the mobility of individual VGCCs in *unc-10(md1117)* mutants using CALM. The two diffusive behaviors of VGCCs were still observed in these mutants and their relative fractions remained unchanged compared to WT (Fig. 2f), suggesting that UNC-10 does not impact the ability of VGGCs to diffuse inside or outside nanodomains. However, VGCCs were significantly slower in *unc-10(md1117)* mutants, with both D_slow_ within nanodomains and D_fast_ across AZ being reduced by half compared to WT (Fig. 2g). This is consistent with a similar decrease in Cav2 diffusion observed after removal of RIM at *Drosophila* NMJs^25^. The channels were also significantly more confined within nanodomains (*unc-10*: 78 ± 23 nm *vs.* WT: 104 ± 23 nm), but their diffusion range across AZ was maintained at WT levels (*unc-10*: 270 ± 28 nm *vs.* WT: 289 ± 29 nm) (Fig. 2h). Together, this indicates that UNC-10 enhances the mobilities of VGCCs both inside and outside nanodomains, in addition to its role as a spatial organizer of VGCC nanodomains at AZs.

### The mobility and spatial organization of nanodomain-associated VGCCs are regulated by UNC-10/RIM mediated coupling to synaptic vesicles

To understand how UNC-10, as a cytomatrix protein, enhances the mobility of VGCCs, we investigated whether the channel diffusion reflects their physical coupling to SVs through UNC-10. As UNC-10 docks SVs by binding to vesicular RAB-3^12,48^ (Fig. 3a), we examined if the mobility of VGCCs would be impacted in *rab-3(js49)* mutants^48^, which have SV docking defect similar to that of *unc-10(md1117)* animals^49,50^. *Rab-3(js49)* mutants exhibited reduced neurotransmission^50^, showing increased resistance to aldicarb and impaired locomotion (Fig. 3b and 3c). However, the loss of RAB-3 had no significant influence on the diffusion coefficients of VGCCs, suggesting that docking defects alone do not account for the decreased channel mobilities observed in *unc-10(md1117)* mutants (Fig. 3d and 3e; Fig. S5a). UNC-10 also binds the priming factor UNC-13L/Munc-13, which physically bridges SVs with the presynaptic membrane^51,52^. We thus examined the channel mobilities in *unc-10(nu487)* animals carrying K77E/K79E mutations, which disrupt interactions between the ZnF domain of UNC-10 and UNC-13L^11,12,17^ (Fig. 3a). *Unc-10(nu487)* mutants were reported to have reduced release probability^17^ and they showed mild uncoordinated phenotypes with near-normal aldicarb-resistance (Fig. 3b and 3c). Again, the diffusion coefficients of VGCCs in these mutants remained similar to WT (Fig. 3d and 3e; Fig. S5b), indicating that dissociating UNC-10 from UNC-13L is not sufficient to reduce the channel mobilities.

**Figure 3.**
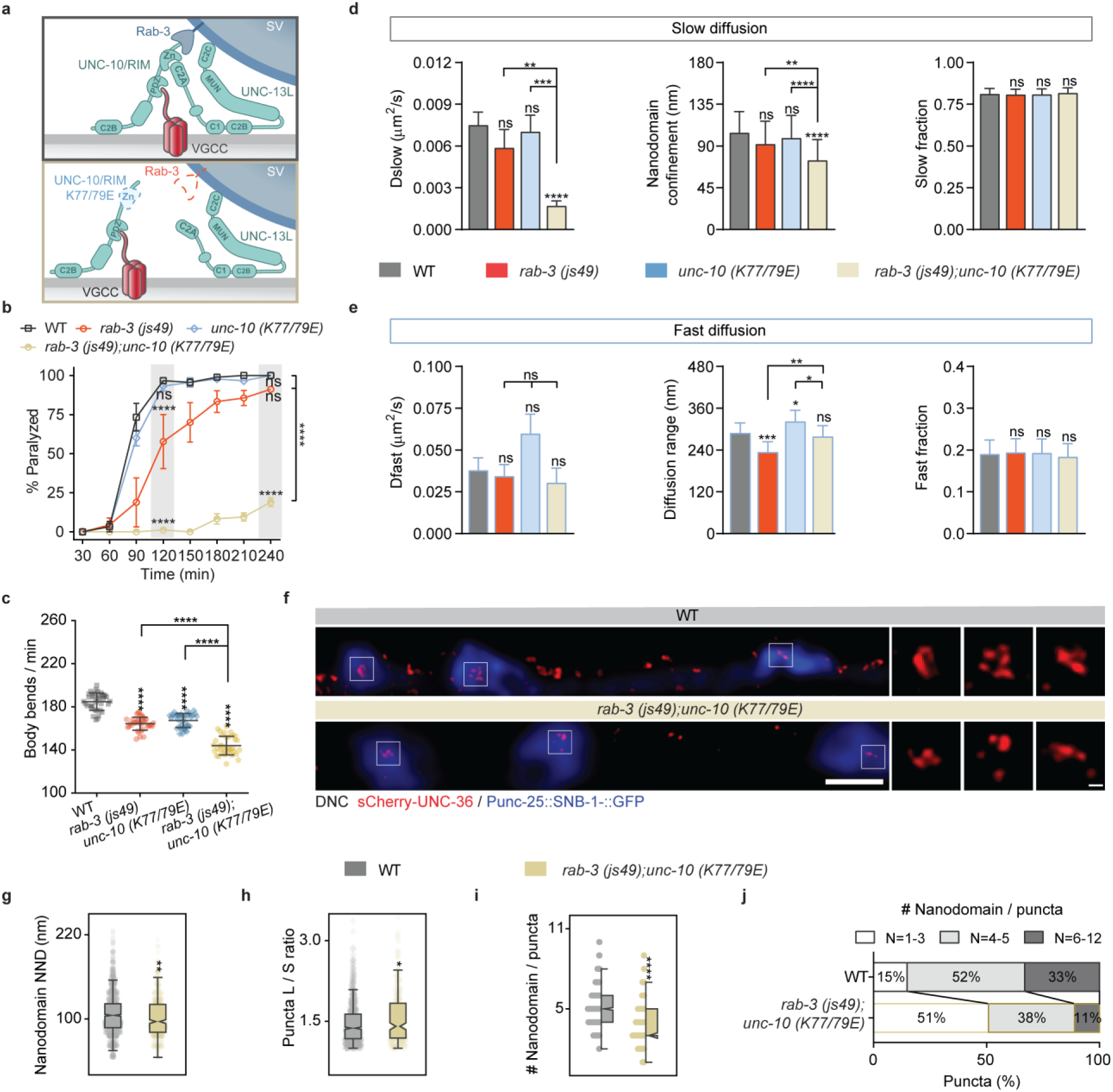
The mobility and spatial organization of nanodomain-associated VGCCs is regulated by UNC-10/RIM mediated SV coupling. **(a)** Schematics of SV coupling to VGCC via UNC-10, RAB-3 and UNC-13L in WT (top) and in *rab-3;unc-10(K77E/K79E)* double mutant (bottom). **(b)** Numbers of body bending per min (mean ± SD) in trashing assays for animals (n = 36) of indicated genotypes. Comparisons by Kruskal-Wallis and Dunn’s test, ****p < 0.0001. **(c)** Percentage of paralyzed animals over time (mean ± SEM) in aldicarb-sensitivity assays, for the indicated genotypes (n = 90). Comparisons to WT at time points by two-way ANOVA and Tukey’s tests, ****p<0.0001; ns, not significant. **(d-e)** Mobility of VGCCs in WT (gray; n = 42,391 trajectories; 58 animals), *rab-3(js49)* (red; n = 33,175 trajectories; 107 animals), *unc-10 (K77/79E)* (blue; n = 29,278 trajectories; 26 animals) and *rab-3; unc-10(K77E/K79E)* mutants (brown; n = 24,455 trajectories; 19 animals), including diffusion coefficients (mean ± SEM), nanodomain confinement size or diffusion range (mean diameter ± localization precision) and fractions (mean ± SEM) for the channels’ slow (d) and fast (e) diffusive behaviors. Comparisons by two-tailed T-test. ****p<0.0001; ***p<0.001; **p<0.01; *p<0.05; ns, not significant. **(f)** Representative STED images of immunostained surface sCherry-UNC-36 (red) overlayed with confocal images of Punc-25::SNB-1::GFP (blue) at the dorsal nerve cords of WT or *rab-3; unc-10(K77E/K79E)* animals. Scale bars, 1 µm and 100 nm for insets. **(g-j)** Quantifications of UNC-36 nanodomain nearest-neighbor distance NND (g), puncta long/short axe ratio (h), nanodomain numbers per puncta (i) and their distribution (j) in STED images from WT (gray; n = 1416 NND; 1076 puncta; 10 animals) and *rab-3; unc-10(K77E/K79E)* mutants (brown; n = 509 NND; 244 puncta; 8 animals). Notched boxes: median ± interquartile range (IQR); whiskers: 1.5xIQR. Comparisons to WT by Kruskal-Wallis and Dunn’s test, ****p < 0.0001; **p<0.01; *p<0.05.

Interestingly, when both interactions of UNC-10 with RAB-3 and UNC-13L were simultaneously disrupted in *rab-3;unc-10(nu487)* double mutants (Fig. 3a), VGCCs within nanodomains exhibited significantly reduced diffusion coefficients (D_slow_) and increased confinement (double mutant: 74 ± 23 nm *vs.* WT: 104 ± 23 nm), phenocopying changes observed in *unc-10(md1117)* animals (Fig. 3d; Fig. S5c). On the contrary, D_fast_ for VGCCs outside nanodomains remained at WT levels (Fig. 3e). This indicates that the two diffusive behaviors of VGCCs are regulated by UNC-10 via distinct mechanisms, with the channel mobility in nanodomains being driven by UNC13L/RAB-3-dependent SV coupling but being independent of SV coupling or of UNC13L/RAB-3 in the rest of the AZ. We note that the larger diffusion range of VGCCs exploring the AZ was differentially impacted in the two single mutants, as it significantly decreased in the absence of RAB-3 (*rab-3*: 234 ± 30 nm *vs.* WT: 289 ± 29 nm), but increased in *unc-10(nu487)* mutants (*unc-10*: 322 ± 32 nm vs. WT: 289 ± 29 nm). These opposing effects offset each other in *rab-3;unc-10(nu487)* double mutants, where the diffusion range remained similar to WT (Fig. 3e). This suggests that the AZ area explored by VGCCs outside nanodomains is promoted by UNC-10/RAB-3-mediated SV docking while it is restricted by UNC-10/UNC-13L interactions. In *rab-3;unc-10(nu487)* double mutants, the neurotransmission and locomotion defects were significantly more severe than in both single mutants, suggesting that the interactions of UNC-10 with RAB-3 and UNC-13L are functionally complementary (Fig. 3b and 3c). Together, these findings show that the diffusion of nanodomain-associated VGCCs is driven by a tripartite UNC-10/RAB-3/UNC-13L SV coupling complex, which is essential for efficient synaptic release.

To test whether this SV coupling complex also participates in organizing VGCC nanodomains, we examined the distribution of VGCCs in *rab-3;unc-10(nu487)* double mutants by STED. We found that the distribution of VGCC nanodomains became more compact and elongated than in WT animals, with their NND and puncta shape (long/short axes ratio) being significantly altered (Fig. 3f-3h). The number of nanodomain per puncta was also significantly reduced, yet it was not impacted to the extent observed in the absence of UNC-10 (Fig. 3i and 3j). This indicates that SV coupling mediated by UNC-10 partially promotes the accumulations of VGCC nanodomains and shapes their spatial organization at AZs.

### The mobility and AZ organization of VGCCs is governed by synaptic vesicle priming and SNARE complex assembly

As UNC-10/UNC-13L interactions couple VGCCs to readily releasable SVs^17^, we next investigated whether the channel mobility is regulated by SV priming. We examined the diffusion of VGCCs in *unc-13(s69L-S-)* null mutants, where the absence of both UNC-13 isoforms abolishes SV priming and synaptic release^51,53^ (Fig. 4a-c; Fig. S6a). In *unc-13(s69L-S-)* animals, nanodomain-associated VGCCs exhibited significantly reduced D_slow_ and increased confinement (*unc-13^L-S-^*: 81 ± 21 nm *vs.* WT: 104 ± 23 nm), phenocopying changes observed in *rab-3;unc-10(nu487)* double mutants (Fig. 4d; Fig. S6b). Because disrupting the interaction of UNC-13L with UNC-10 in *unc-10(nu487)* animals did not influence the channel mobility, these results suggest that VGCC diffusion in nanodomains is impacted by SV priming via both UNC-13L and UNC-13S rather than by simple interaction between UNC-13L and UNC-10. To further test this, we studied the diffusion of VGCCs in *unc-13(e51L-S+)* mutants, where priming levels are slightly restored by endogenous expression of only the UNC-13S isoform, which lacks the UNC-10-binding domain, but contains the core regions required for priming^53^ (Fig. 4a). In *unc-13(e51L-S+)* animals, neurotransmission was significantly improved compared to *unc-13(s69L-S-)* mutants (Fig. 4c; Fig. S6a), but it remained defective compared to WT, consistent with UNC-13L being critical for efficient SV release^17,53^. When expressing the UNC-13S isoform, both D_slow_ and the confinement size of nanodomain-associated VGCCs were partially restored, with D_slow_ remaining slower than in WT, albeit short of statistical significance (p = 0.219, Fig. 4d; Fig. S6c and S6d). This indicates that moderately improved SV priming levels by UNC13S expression is sufficient to promote the mobility of slow VGCCs, but that UNC13L-mediated priming is required to restore WT-level diffusion in AZ nanodomains. VGCCs outside nanodomains were impacted in similar ways. D_fast_ decreased in *unc-13(s69L-S-)* mutants, albeit short of statistical significance (p = 0.178), and it was only partially restored to WT-levels in *unc-13(e51L-S+)* animals (Fig. 4e). Compared to WT, the channel diffusion range became significantly smaller when both UNC-13 isoforms were ablated (*unc-13^L-S-^*: 240 ± 23 nm *vs.* WT: 289 ± 29 nm) but it was restored by the sole expression of UNC-13S (*unc-13^L-S+^*: 310 ± 28 nm *vs.* WT: 289 ± 29 nm; Fig. 4e). Together, these results indicate that UNC-13-mediated SV priming modulates the mobility of VGCCs both within and outside nanodomains at the AZ.

**Figure 4.**
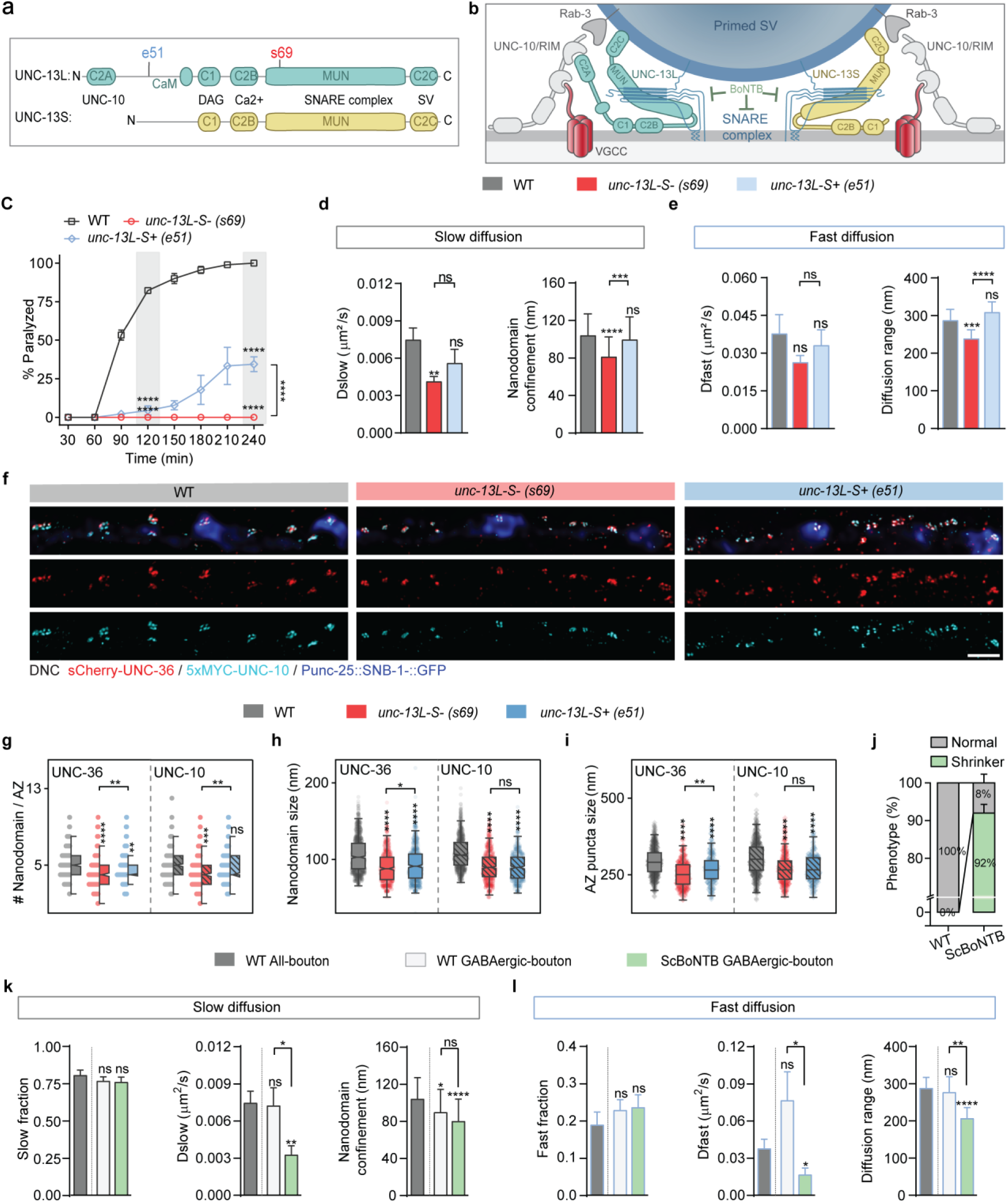
The mobility and AZ organization of VGCCs is governed by SV priming and SNARE complex assembly. **(a)** Domain maps and binding parters of UNC-13 isoforms. **(b)** Schematic of SV priming by UNC-13L (turquoise) and UNC-13S (yellow) isoforms. **(c)** Percentage of paralyzed animals over time (mean ± SEM) in aldicarb-sensitivity assays, for the indicated genotypes (n = 90). Comparisons to WT at time points or across genotypes at t = 240 min by two-way ANOVA and Tukey’s tests, ****p<0.0001. **(d-e)** Mobility of VGCCs in WT (gray; n = 42,391 trajectories; 58 animals), *unc-13L-S- (s69)* (red; n = 31,355 trajectories; 35 animals) and *unc-13L-S+ (e51)* mutants (blue; n = 19,899 trajectories; 34 animals), including diffusion coefficients (mean ± SEM) and nanodomain confinement size or diffusion range (mean diameter ± localization precision) for the channels’ slow (d) and fast (e) diffusive behaviors. Comparisons by two-tailed T-test. ****p<0.0001; ***p<0.001; **p<0.01; *p<0.05; ns, not significant. **(f)** Representative dual-color STED images of immunostained surface sCherry-UNC-36 (red) and 5xMYC-UNC-10 (cyan) overlayed with confocal images of Punc-25::SNB-1::GFP (blue) at the dorsal nerve cords of WT, *unc-13L-S- (s69)* and *unc-13L-S+ (e51)* animals. Scale bars, 1 µm. **(g-i)** Quantifications of UNC-36 and UNC-10 nanodomain numbers per AZ (g), nanodomain sizes (h), and puncta size at AZ (i) in WT (gray; n = 516 AZ puncta; 993 UNC-36 and 786 UNC-10 nanodomains; 10 animals), *unc-13L-S- (s69)* (red; n = 320 AZ puncta; 607 UNC-36 and 641 UNC-10 nanodomains; 6 animals) and *unc-13L-S+ (e51)* mutants (blue; n = 205 AZ puncta; 655 UNC-36 and 473 UNC-10 nanodomains; 4 animals). Notched boxes: median ± interquartile range (IQR); whiskers: 1.5xIQR. Comparisons by Kruskal-Wallis and Dunn’s test, ****p < 0.0001; **p<0.01; *p<0.05; ns, not significant. **(j)** Percentage of animals displaying normal (grey) or “shrinker” (green) locomotory phenotypes (mean ± SEM) in WT and transgenic animals expressing ScBoNTB in GABAergic synapses (n = 50), over 3 independent batches of animals. **(k-l)** Mobility of VGCCs in WT animals at all boutons (dark gray; n = 42,391 trajectories; 58 animals), in WT animals at GABAergic boutons only (light gray; n = 19,034 trajectories; 42 animals) and in ScBoNTB-expressing animals at GABAergic boutons only (green; n = 11,549 trajectories; 36 animals), including fractions (mean ± SEM), diffusion coefficients (mean ± SEM), and nanodomain confinement size or diffusion range (mean diameter ± localization precision) for the channels’ slow (k) and fast (l) diffusive behaviors. Comparisons by two-tailed T-test. ****p<0.0001; ***p<0.001; **p<0.01; *p<0.05; ns, not significant.

To investigate if the structural organization of VGCCs at AZs was also modulated by UNC-13-mediated SV priming, we imaged VGCCs and UNC-10 by STED in the two *unc-13* mutants. For both VGCCs and UNC-10, the number of nanodomains per AZ and the size of nanodomains were significantly lower in *unc-13(s69L-S-)* mutants compared to WT (Fig. 4f-4h; Fig. S6f), with the distribution of UNC-10 nanodomains being more compact, but that of VGCC nanodomains being unchanged (NND, Fig. S6e). The mean AZ puncta size of VGCCs and UNC-10 were both significantly reduced (AED, Fig. 4i), indicating that AZs were globally smaller when UNC-13 mediated SV priming was abolished, which led to a decrease in the diffusion range of fast VGCCs. With the expression of UNC-13S in *unc-13(e51L-S+)* mutants, VGCC nanodomain numbers, their size and the AZ puncta sizes were only partially restored compared to WT animals (Fig. 4f-4i; Fig. S6f). This indicates that UNC-13S-mediated SV priming influences the spatial distribution of VGCCs, but that UNC-13L-mediated priming is required to maintain a WT structural organization of the channels at AZs, consistent with our diffusion measurements. For UNC-10, the sole expression of UNC-13S fully restored the number of nanodomains per AZ and their NND to WT levels (Fig. 4g; Fig. S6e). Yet, in the absence of UNC-13L, the nanodomain size of UNC-10 and its AZ puncta size remained well below WT levels and non-significantly different from *unc-13(s69L-S-)* mutants (Fig. 4h and 4i). These results indicate that the number and distribution of UNC-10 nanodomains are also modulated by UNC-13S-mediated SV priming, while UNC13-L mediated priming contributes to the size of individual UNC-10 nanodomains and that of the AZ. Surprisingly, the apparent rescue of VGCC nanodomain confinement and AZ diffusion range to WT levels in *unc-13(e51L-S+)* animals (Fig. 4e) did not fully correlate with the nanodomain and AZ puncta sizes for UNC-10 in those mutants (Fig. 4i), suggesting that, in the absence of UNC-13L, UNC-13S-mediated SV priming induces VGCCs to diffuse beyond spatial boundaries defined by UNC-10.

We next investigated whether the mobility of VGCCs is modulated by SNARE complex assembly during SV priming and whether SV coupling, at this molecular step, impacts the channels diffusion. We utilized the transgenic expression of the botulinum neurotoxin serotype B light chain protease (ScBoNTB), which blocks full SNARE complex assembly by cleaving the v-SNARE vesicular SNB-1^54^ (Fig. 4b). As previously reported, expressing ScBoNTB in GABAergic neurons silenced inhibitory neurotransmissions^54^ and 92% of transgenic animals showed “shrinker” phenotypes (Fig.4j). For proper comparison, we first reanalyzed the mobility of VGCCs specifically at GABAergic synapses in WT animals by selecting trajectories only located at the tip of GABAergic boutons (Fig. S6g). GABAergic VGCCs also exhibited two diffusive behaviors, with respective fractions and diffusion coefficients that were non-significantly different from those observed across all buttons, although D_fast_ appeared slightly faster (Fig. 4k and 4l; Fig. S6h). Only the confinement size of nanodomain-associated GABAergic VGCCs was smaller (WT^GABAergic^: 90 ± 25 nm *vs.* WT^All^: 104 ± 23 nm) (Fig. 4k). We then tracked GABAergic VGCCs in ScBoNTB-expressing animals. We found that both D_slow_ and D_fast_ significantly decreased, an indication that the mobility of VGCCs inside and outside AZ nanodomains is indeed modulated by a molecular priming of SVs that requires full SNARE assembly (Fig. 4k and 4l; Fig. S6i). The diffusion range of VGCCs outside nanodomains was also significantly reduced (ScBoNTB^GABAergic^: 207 ± 29 nm *vs.* WT^GABAergic^: 278 ± 42 nm; Fig. 4j). But the confinement size of nanodomain-associated VGCCs was unchanged compared to GABAergic WT (Fig. 4k), suggesting that it is independent of SNARE complex assembly. Altogether, these results demonstrate that, downstream of UNC-10 mediated channel-vesicle coupling and as part of UNC-13 mediated priming, the complete assembly of SNARE complexes that clamp SVs is a critical step that modulates VGCC diffusions at AZs.

### The two mobilities of VGCCs are differentially modulated by UNC-13 isoforms and synaptic vesicle priming levels

We then examined if the mobility of VGCCs would positively scale with SV priming levels by tuning the balance between UNC-13 isoforms and the t-SNAREs inhibitor TOM-1/Tomosyn. In *C. elegans*, removal of TOM-1 markedly upregulates the pool of primed SVs throughout synaptic terminals^55,56^, and also partially rescues synaptic release defects in *unc-13* mutants^53^. We therefore compared the diffusion of VGCCs in *tom-1;unc-13^L-S-^* versus *tom-1;unc-13^L-S+^*double mutants, where SV priming levels are progressively restored, and in *tom-1(nu468)* single mutants where priming levels are largely augmented^53,56^. As previously reported, both double mutants exhibited significantly improved locomotion and neurotransmissions compared to the corresponding *unc-13* single mutants, with SV release rescue being stronger in the presence of UNC-13S (Fig. 5a; Fig. S7a). We found that D_slow_ for nanodomain-associated VGCCs was partially restored in *tom-1;unc-13 ^L-S-^* animals, while their confinement size fully recovered to WT-level (*tom-1;unc-13^L-S-^*: 111 ± 22 nm *vs.* WT: 104 ± 23 nm, Fig.5b), confirming that slow-diffusing VGCCs are modulated by SV priming. However, despite enhanced SV priming levels, the mobility of VGCCs inside nanodomains was not further improved by the expression of UNC-13S in *tom-1;unc-13^L-S+^* double mutants, as it remained similar to *tom-1;unc-13^L-S-^* animals and to *unc-13(e51L-S+)* single mutants (Fig. 5b; Fig. S7b-S7f). These results suggest that UNC-13S and TOM-1 have limited effect on the mobility of nanodomain-associated VGCCs, which are primarily regulated by UNC-13L. Indeed, the mobility of VGCCs in nanodomains did not further increase with augmented SV priming levels in *tom-1(nu468)* single mutants (Fig. 5a and 5b; Fig. S7e).

**Figure 5.**
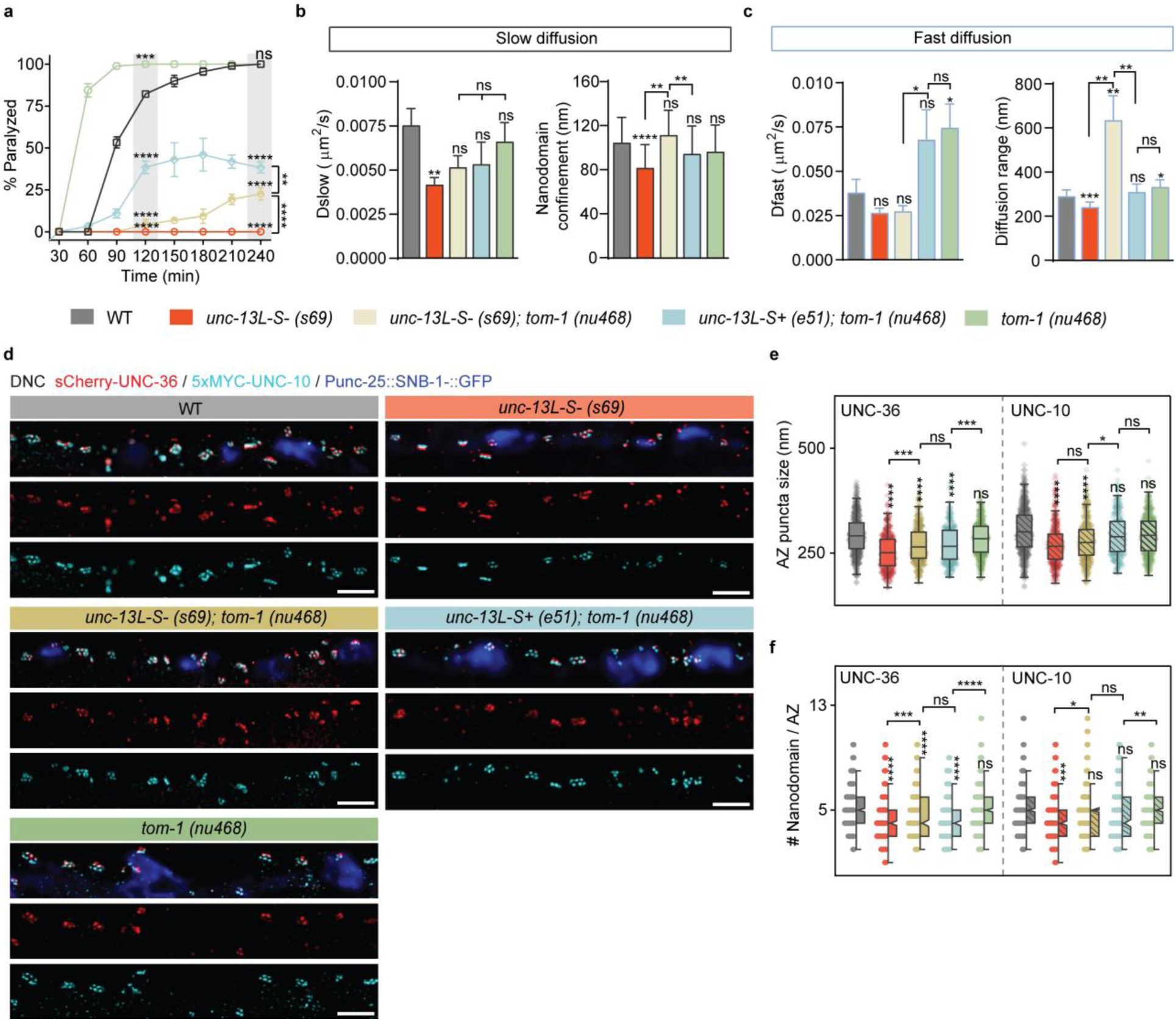
Distinct mobilities and nanodomain organization of VGCCs are differentially modulated by SV priming levels. **(a)** Percentage of paralyzed animals over time (mean ± SEM) in aldicarb-sensitivity assays, for the indicated genotypes (n = 90). Comparisons to WT at time points t = 120min and across genotypes at t = 240 min by two-way ANOVA and Tukey’s tests, ****p<0.0001; ***p<0.001; **p<0.01. **(b-c)** Mobility of VGCCs in WT (gray; n = 42,391 trajectories; 58 animals), *unc-13L-S- (s69)* (red; n = 31,355 trajectories; 35 animals), *unc-13L-S- (s69);tom-1(nu468)* (brown; n = 25,309 trajectories; 15 animals)*, unc-13L-S+ (e51);tom-1(nu468)* mutants (cyan; n = 18,963 trajectories; 15 animals), and *tom-1(nu468)* mutants (green; n = 34,062 trajectories; 25 animals), including diffusion coefficients (mean ± SEM) and nanodomain confinement size or diffusion range (mean diameter ± localization precision) for the channels’ slow (b) and fast (c) diffusive behaviors. Comparisons by two-tailed T-test. ***p<0.001; **p<0.01; *p<0.05; ns, not significant. **(d)** Representative dual-color STED images of immunostained surface sCherry-UNC-36 (red) and 5xMYC-UNC-10 (cyan) overlayed with confocal images of Punc-25::SNB-1::GFP (blue) at the dorsal nerve cords of WT, *unc-13L-S-(s69)*, *unc-13L-S-(s69);tom-1(nu468)*, *unc-13L-S+ (e51);tom-1(nu468)* and *tom-1(nu468)* animals. Scale bars, 1 µm. **(e-f)** Quantifications of UNC-36 and UNC-10 puncta size at AZ (e) and nanodomain numbers per AZ (f) in WT (gray; n = 516 AZ puncta; 10 animals), *unc-13L-S-(s69)* (red; n = 320 AZ puncta; 6 animals), *unc-13L-S- (s69);tom-1(nu468)* (brown; n = 349 AZ puncta; 10 animals)*, unc-13L-S+ (e51);tom-1(nu468)* mutants (cyan; n = 276 AZ puncta; 8 animals) and *tom-1(nu468)* mutants (green; n = 529 AZ puncta; 10 animals). Notched boxes: median ± interquartile range (IQR); whiskers: 1.5xIQR. Comparisons by Kruskal-Wallis and Dunn’s test, ****p < 0.0001; *p<0.05; ns, not significant.

On the contrary, for VGCCs outside nanodomains, both D_fast_ and their diffusion range significantly increased in *tom-1(nu468)* animals (*tom-1*: 331 ± 33 nm *vs.* WT: 289 ± 29 nm; Fig. 5c). This enhanced mobility was suppressed when UNC-13 was ablated in *tom-1;unc-13^L-S-^* double mutants, where D_fast_ was similar to that in *unc-13(s69L-S-)* single mutants (Fig. 5c). However, the increase in D_fast_ was largely preserved by UNC-13S expression in *tom-1;unc-13^L-S+^*double mutants, who displayed significantly faster VGCC diffusion than in *tom-1;unc-13^L-S-^* animals (Fig. 5c; Fig. S7d-S7f). This indicates that the faster mobility of VGCCs outside nanodomains in *tom-1(nu468)* mutants is primarily UNC-13S-dependent, conforming with a previously reported upregulation of UNC-13S abundance in these animals^56^. Together, these findings show that the mobility of VGCCs is predominantly governed by UNC-13L-mediated SV priming inside nanodomains, and that it scales with a pool of TOM-1 regulated and UNC-13S-primed SVs across the rest of the AZ. Interestingly, the diffusion range of VGCCs outside nanodomains was dramatically expanded in *tom-1;unc-13^L-S-^* double mutants (*tom-1;unc-13^L-S-^*: 632 ± 112 nm *vs.* WT: 289 ± 29 nm; Fig. 5c), in stark contrast with the reduction observed in *unc-13(s69L-S-)* single mutants. As the number of membrane-contacting SVs located at the AZ periphery (400-600 nm away from the DPs) substantially increases in *tom-1;unc-13^L-S-^*animals^55^, our observation suggests that VGCCs can diffuse far across the synaptic area where UNC-13-independent SV priming is upregulated. Concurrently, in the same *tom-1;unc-13^L-S-^* animals, the size of UNC-10 puncta was significantly reduced (Fig. 5e), indicating that, in the absence of UNC-13, the diffusion range of fast VGCCs is set by the location of primed SVs rather than by the distribution of UNC-10 at AZ puncta. When SV priming was controlled by UNC-13S in *tom-1;unc-13 ^L-S+^* double mutants, the diffusion range of fast VGCCs was restored to WT levels (*tom-1;unc-13^L-S+^*: 307 ± 37 nm *vs.* WT: 289 ± 29 nm), together with the median puncta size of UNC-10 (Fig. 5c-5e). And when UNC-13S-mediated SV priming was upregulated in *tom-1(nu468)* animals, this same diffusion range significantly increased compared to WT (*tom-1*: 331 ± 33 nm *vs.* WT: 289 ± 29 nm; Fig. 5c), again exceeding the median UNC-10 puncta size (Fig. 5c-5e). Together, these observations indicate that outside nanodomains, the mobility of VGCCs is effectively governed by UNC-13S-mediated SV priming, and that the channel diffusion range can expand when UNC-13-independent and spatially imprecise SV priming is upregulated.

We also examined if increasing priming levels by TOM-1 removal modulates the AZ organization of VGCC and UNC-10 nanodomains. We found that the defective nanodomain numbers and puncta size of VGCCs we had observed when ablating the two UNC-13 isoforms were partially restored by simultaneous removal of TOM-1 in *tom-1;unc-13^L-S-^* double mutants, although they both remained significantly lower than WT (Fig. 5d-5f). The median number and NND of UNC-10 nanodomains were back to WT-level, yet UNC-10 puncta sizes did not recover (Fig. 5d-5f; Fig. S7g-S7i). These results are similar to those found in *unc-13(e51L-S+)* mutants, where SV priming levels are improved and comparable to those of *tom-1;unc-13^L-S-^* animals^57^. Further improvements of SV priming in *tom-1;unc-13 ^L-S+^* double mutants fully restored UNC-10 puncta sizes to WT levels, but the organization of VGCC nanodomains remained partially defective (Fig. 5d-f), confirming that UNC-13L is important for organizing VGCCs at the AZ. In *tom-1(nu468)* animals, where both UNC-13 isoforms are present and UNC-13S-mediated priming is predominantly upregulated, the spatial organizations of both VGCC and UNC-10 nanodomains were indistinguishable from WT (Fig. 5d-5f). This suggests that VGCC and UNC-10 organizations scale with SV priming levels up to a limit set by UNC-13L-activity. Collectively, these findings demonstrate that UNC-13 mediated SV priming modulates the organization of VGCC and UNC-10 nanodomains in a parallel and progressive manner. Notably the increase in UNC-10 nanodomain numbers is followed by an expansion of AZ puncta sizes, and the organization of VGCC nanodomains exhibits greater sensitivity to priming levels than that of UNC-10.

### Primed synaptic vesicles can modulate the mobilities and multi-nanodomain organization of VGCCs downstream of UNC-10

As the diffusive behaviors of VGCCs are promoted by UNC-10 and SV priming, we tested whether they both modulate VGCC mobility in the same molecular pathway by examining the channels diffusion in *tom-1;unc-10* double mutants. Interestingly, the defective D_slow_ and confinement sizes of nanodomain-associated VGCCs we observed in *unc-10(md1117)* single mutants were fully restored to WT-level in *tom-1;unc-10* double mutants (Fig. 6a; Fig. S8a). Likewise, the diffusion coefficient of VGCCs outside nanodomains was also significantly increased, with D_fast_ being faster than in WT, albeit short of significance (p = 0.193, Fig. 6b), and being slower than the augmented D_fast_ observed in *tom-1(nu468) single* mutants. Thus, upregulating SV priming levels can partially bypass the role of UNC-10 in modulating the mobility of VGCCs. The diffusion range of fast VGCCs in *tom-1;unc-10* double mutants showed significant increase compared to WT (*unc-10*;*tom-1*: 336 ± 27 nm *vs.* WT: 289 ± 29 nm; Fig. 6b; Fig. S8a), phenocopying changes observed in *tom-1(nu468)* animals and confirming that the diffusion range is set by primed SVs independently of UNC-10. Together, it demonstrates that UNC-10 functions to couple mobile VGCCs to primed SVs, which can directly modulate the channel mobilities downstream of UNC-10. TOM-1 removal also largely rescued the impaired neurotransmissions in *unc-10(md1117)* background (Fig. 6c) as previously reported^57^. Since UNC-10 is not required for molecular priming^17,58^, the rescue of synaptic releases through TOM-1 ablation in double mutants likely stems from the upregulated SV priming levels and the restored VGCC mobility, which might compensate for reduced SV docking^45^.

**Figure 6.**
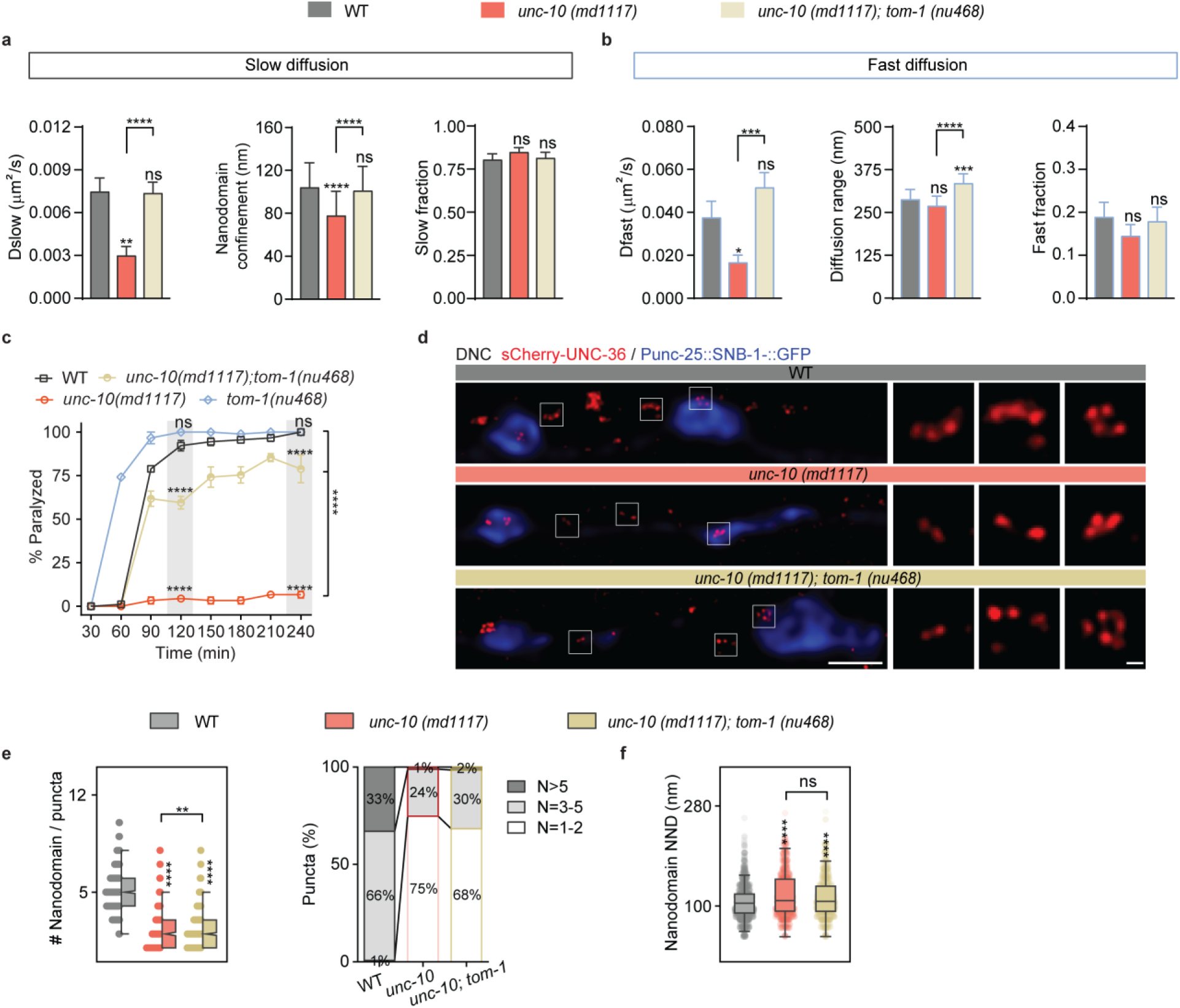
Primed SVs modulate the mobilities and multi-nanodomain organization of VGCCs downstream of UNC-10. **(a-b)** Mobility of VGCCs in WT (gray; n= 42,391 trajectories; 58 animals); *unc-10 (md1117)* (red; n= 18,645 trajectories; 34 animals) and *unc-10(md1117);tom-1(nu468)* (brown; n = 20,882 trajectories; 26 animals) mutants, including diffusion coefficients (mean ± SEM), nanodomain confinement size or diffusion range (mean diameter ± localization precision) and fractions (mean ± SEM) for the channels’ slow (a) and fast (b) diffusive behaviors. Comparisons by two-tailed T-test. ****p < 0.0001; ***p<0.001; **p<0.01; *p<0.05; ns, not significant. **(c)** Percentage of paralyzed animals over time (mean ± SEM) in aldicarb-sensitivity assays, for the indicated genotypes (n = 90). Comparisons to WT at time points or across genotypes at t = 240 min by two-way ANOVA and Tukey’s tests, ****p<0.0001; ns, not significant. **(d)** Representative STED images of immunostained surface sCherry-UNC-36 (red) overlayed with confocal images of Punc-25::SNB-1::GFP (blue) at the dorsal nerve cords of WT, *unc-10 (md1117)* and *unc-10(md1117);tom-1(nu468)* animals. Scale bars, 1 µm and 100 nm for insets. **(e-f)** Quantifications of UNC-36 nanodomain numbers per puncta (e; left), distribution (e; right) and nearest-neighbor distance NND (f) in WT (gray; n = 1076 puncta; 1416 NND; 10 animals), *unc-10 (md1117)* (red; n = 634 puncta; 855 NND; 7 animals) and *unc-10(md1117);tom-1(nu468)* (brown; n = 1087 puncta; 748 NND; 8 animals) mutants. Notched boxes: median ± interquartile range (IQR); whiskers: 1.5xIQR. Comparisons by Kruskal-Wallis and Dunn’s test, ****p < 0.0001; **p<0.01; ns, not significant.

Finally, we examined whether increasing priming levels can rescue the defective organizations of VGCC nanodomains in the absence of UNC-10. In *tom-1;unc-10* animals, the number of VGCC nanodomains per puncta was slightly but significantly increased compared to *unc-10(md1117)* single mutants, yet it remained largely reduced compared to WT (Fig. 6d and 6e). In contrast, the NND between nanodomains was non-significantly different from that in *unc-10(md1117)* animals, showing a more dispersed organization compared to WT (Fig. 6f). These results demonstrate that the multi-nanodomain organization of VGCCs at AZ requires UNC-10 and indicate that SV priming levels regulate the spatial distribution of VGCC nanodomains through UNC-10.

## Discussion

Using single channel tracking and super-resolution imaging in *C. elegans*, we dissected how mobile VGCCs are spatiotemporally aligned with synaptic vesicles for precise neurotransmission. We discovered that individual N-type VGCCs adopt distinct mobilities determined by their spatial distribution and the molecular machinery coupling them to readily releasable SVs (Fig. 7). VGCCs tethered to primed SVs via a tripartite UNC-10/RAB-3/UNC-13L complex diffuse slowly in multiple sub-AZ nanodomains that align with UNC-10/RIM scaffolds, whereas VGCCs coupled to UNC-13S-primed SVs move significantly faster across the larger AZ puncta. The spatially correlated multi-nanodomain arrangement of VGCCs and UNC-10 closely resembles Cav2 and RIM1/2 nanoclusters observed at rodent hippocampal synapses^52,59^, consistent with nematode synapses exhibiting a multi-release-site organization similar to mammals^60^. Heterogeneity in nanodomain number and size across synapses further points to a modular intra-AZ organization that may enable local tuning of synaptic strength. Together, this nanoscale organization and the distinct diffusion regimes of VGCCs likely give rise to diverse channel-vesicle coupling modes that shape SV release probability.

**Figure 7.**
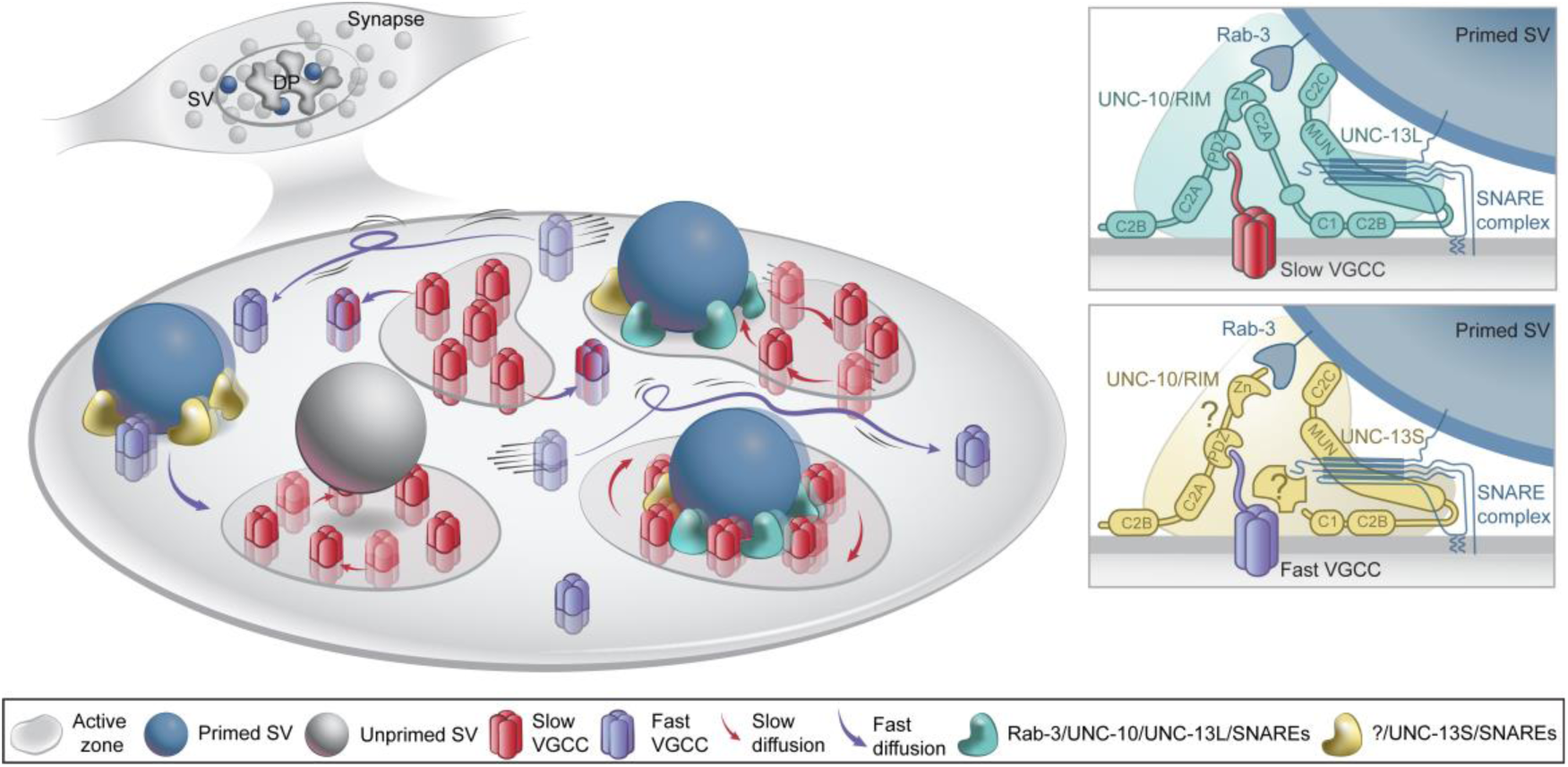
Schematic illustrating the heterogeneous mobility and multi-nanodomain organization of VGCCs at the AZ. In the AZ surrounding the dense projection (DP, top left), VGCCs (red) exhibit slow diffusion within multiple ∼100 nm nanodomains and transition to a faster diffusing mode (purple) outside these regions while remaining confined within a ∼300 nm AZ area. Within nanodomains, primed synaptic vesicles (SVs, blue) drive VGCC mobility via molecular coupling mediated by UNC-10/RAB-3/UNC-13L/SNARE complexes (turquoise; top right) whereas across the broader AZ, VGCC dynamics are modulated by UNC-13S/SNARE complexes together with additional factors (yellow; bottom right).

In this context, UNC-10 emerges as a key regulator of VGCC mobility and nanodomain organization. We found that UNC-10 promotes VGCC diffusion by coupling channels to primed SVs, and that priming levels modulate both channel dynamics and multi-nanodomain organization. As an SV docking factor and a direct modulator of VGCCs, UNC-10/RIM drives channel accumulation at AZs^7,9,47^, stabilizing them and securing their coupling distances to vesicles^8,18,61^. Consistent with these functions, UNC-10 facilitates the accumulation and tight organization of multiple VGCC nanodomains at AZs. However, VGCCs can form nanodomains independently of UNC-10 and, counterintuitively, UNC-10 enhances channel mobility. Thus, rather than simply stabilizing VGCCs, UNC-10 modulates channel dynamics and shapes their nanotopography. The influence of UNC-10 on VGCC diffusion appears conserved across species, as Ca_V_2.1 isoforms harboring or lacking RIM1/2 binding motifs exhibit different mobilities at rodent hippocampal synapses^23^, and Cac/Ca_V_2 displays similarly slower diffusion in the absence of RIM at *Drosophila* NMJs^25^.

Importantly, at *C. elegans* NMJs, VGCCs within sub-AZ nanodomains and across the broader AZ exhibit two different mobility regimes that both arise from coupling to primed SVs but are governed by distinct molecular mechanisms. To dissect these mechanisms, we altered vesicle priming through UNC-13 isoforms, the t-SNARE inhibitor TOM-1, and SNARE complex blockade, and found that channel dynamics depend on UNC-13-mediated vesicle priming following full SNARE assembly and SV-plasma membrane bridging. Within nanodomains, VGCC dynamics are primarily driven by SV priming mediated through UNC-10/RAB-3/UNC-13L complexes, with additional contributions from the UNC-13S isoform. Outside nanodomains, however, this tripartite complex does not regulate VGCC mobility but instead influences the channel distribution across AZ. In this region, VGCC dynamics scale with pools of UNC-13S-primed or UNC-13-independent SVs, which are physiologically inhibited by TOM-1. Accordingly, blocking SNARE assembly with ScBoNTB significantly reduces both VGCC diffusion and restricts their AZ distribution, whereas removal of TOM-1 partially restores defective channel dynamics in the absence of UNC-13 or even UNC-10. Thus, UNC-10-mediated SV coupling drives VGCC diffusion following SNARE assembly, while primed SVs can further modulate VGCC behavior beyond UNC-10-dependent coupling. These findings establish a causal link between SV priming levels and channel dynamics and may mechanistically explain why conditions that impair priming and SNARE assembly, such as intracellular Ca²⁺ chelation or silencing of synaptic activity, are associated with reduced VGCC mobility but increased Syntaxin-1A/t-SNARE diffusion^23,24,62^.

Accordingly, the AZ distribution of fast VGCCs is primarily dictated by the location of primed SVs rather than the multi-nanodomain organization of UNC-10. This is evident when simultaneous removal of UNC-13 and TOM-1 leads to an expansion of VGCC diffusion range toward distal AZ regions, where UNC-13-independent SV priming is upregulated^55^. The modest expansion observed after disrupting UNC-10/UNC-13L interactions likely reflects a broadening in primed SV distribution, with UNC-13L becoming less anchored at the AZ^57^. Conversely, removing RAB-3 significantly reduces the diffusion range of VGCCs, as impaired SV docking limits their spatial distribution across AZ. These observations indicate that VGCC topography is dynamically instructed by the spatial distribution of primed SVs rather than solely imposed by AZ scaffolds. Consistent with this relationship, SV priming levels progressively regulate the number and size of VGCC and UNC-10 nanodomains, with the organization of VGCCs being more sensitive to priming levels. These results complement previous findings that Ca_V_2 clustering and vesicle priming represent two distinct sub-assemblies tethered together by RIM^63^. Our data further indicates that this tethering is itself regulated in a priming-level-dependent manner, with primed vesicles reciprocally regulating UNC-10/RIM and directly shaping VGCC diffusion.

Collectively, our findings support a model in which primed SVs actively drive VGCC diffusion through physical coupling, coordinating the movement of membrane-embedded channels with that of adjacent primed SVs^64–68^. Such coordination likely secures the coupling distance between the VGCC calcium donors and the SV-tethered synaptotagmin calcium sensors^1,69,70^ thereby ensuring precise neurotransmission within the highly dynamic environment of the AZ. Furthermore, coupling distinct VGCC mobility to different pools of primed SVs suggests a level of synaptic modularity that supports diverse release properties.

Placing this model in a broader context, studies at *Drosophila* and mammalian synapses^71–75^, together with functional analyses in *C. elegans* indicate that distinct channel-vesicle coupling distances underlie different release properties, with fast release requiring UNC-13L and slow release depending on both UNC-13L and UNC-13S^57,76^. Coupling distance has conventionally been defined as the distance between docked SVs and the DP at the center of the AZ, where DP markers appear as homogeneous puncta in diffraction-limited images^27,47,58^. Our studies reveal a heterogeneous, multi-nanodomain topography of UNC-10 and VGCCs that closely resembles the architecture of DP^44^. This spatial organization, together with differences in VGCC mobility and their coupling to distinct UNC-13 isoforms, supports a multidimensional framework for generating diverse SV release properties. Within this framework, UNC-13L-primed SVs preferentially dock in nanodomains and tether slow-diffusing VGCCs, whose restricted mobility promotes concentrated calcium influx^24^ for fast release. In contrast, UNC-13S-primed SVs dock outside nanodomains or at distal-AZ regions, where they couple to fast-diffusing VGCCs, whose higher mobility yields more diffuse calcium influx for slow release. This framework further implies that differences in channel mobility contribute to defining microdomain- and nanodomain-coupling modes^20^.

Whether and how primed SVs tether fast-diffusing VGCCs outside nanodomains remains unclear. Because fast-diffusing VGCCs are not coupled to the tripartite UNC-10/RAB-3/UNC-13L complex, the assembled SNARE complex itself may represent a candidate. While mammalian Cav2 channels contain a “synprint” motif that directly binds SNARE complexes^4,77^, UNC-2/Cav2 in *C. elegans* lacks such a sequence^78^, and identifying the mechanisms that mediate coupling to fast-diffusing VGCCs will require further investigation. Despite this evolutionary divergence, the principle of vesicle-channel coupling may be more fundamental than the specific protein-protein interactions that mediate it^79^. Another open question is what determines the transition between slow and fast diffusion modes for VGCCs. It will be important to test whether coupled SV-VGCC units move in and out of VGCC nanodomains together, potentially supporting transitions between tightly primed (TS) and loosely primed (LS) states^80^. Addressing these questions will provide further insight into how synapses exploit VGCC dynamics to achieve both reliability and plasticity.

In conclusion, this study identifies a mechanism by which presynaptic VGCC diffusion is actively regulated by the release machinery and the priming status of SVs. The coupling of distinct channel mobilities to different pools of primed vesicles reveals a previously unrecognized layer of AZ organization that fine-tunes neurotransmitter release. Ultimately, this work establishes a mechanistic framework to investigate the role of VGCC dynamics in synaptic function and disease.

## Acknowledgement

We thank Dr. Dion Dickman and Dr. Kaikai He for providing important resource support and advice for STED imaging and analysis. We thank Dr. Hong Zhan for advice on CALM imaging and genome editing of *C. elegans*, and Dr. Carolyn Phillips for resource support on genetics. We also thank Dr. Jean-Louis Bessereau for kindly sharing the unc-25p::snb-1::gfp strain, Dr. Peri Kurshan for her kind gifts of GFP-UNC-2 related strains. We additionally thank Dr. Joshua Kaplan for sharing tom-1(nu468) related strains, Dr. Alexander Gottschalk for kindly providing the unc-47p::eGFP::ScBoNTB strain and Dr. Zhitao Hu for sharing the unc-10 (nu487 K77/79E) strain. We also thank Katya Kadyshevskaya for helping with illustrations in some of the figures. This work was funded by NIH grant NS114911 from the National Institute of Neurological Disorders and Stroke (NINDS). YZ was partially supported by the NIGMS training grant T32GM118289-04 Chemistry Biology Interface Training Grant at the University of Southern California. Some strains were obtained from the Caenorhabditis Genetics Center (CGC) that is funded by the NIH Office of Research Infrastructure Programs (P40 OD010440).

## Methods

### *C. elegans* culture and maintenance

All *Caenorhabditis elegans* nematode strains were raised on nematode growth media (NGM) plates at 20°C and fed on OP50 *Escherichia coli* as previously described^33^. All *C. elegans* used for experiments were adult hermaphrodites. Worms were handled using a Nikon dissecting microscope (SM2745). A full list of strains is available in Supplementary Table 1.

### Molecular biology

For genotyping, worms were lysed in worm lysis buffer as previously described^30^. Worm lysates were amplified using either 2xTaq PCR Premix (Bioland) or Phusion High-fidelity DNA Polymerase (NEB). For CRISPR-Cas9 genome editing, direct microinjection of *in vitro*–assembled CRISPR-Cas9 and single chimeric guide RNA (sgRNA) ribonucleoprotein complexes into the gonads of hermaphrodite worms were performed as previously described^30,81^. Cas9 endonuclease was expressed *in vitro* using DE3 GOLD (Agilent) cells transformed with the nm2973 plasmid^82^ and was prepared to 10 μg/μl. Single-stranded DNA oligonucleotides (ssODNs) were ordered from Integrated DNA Technology (IDT). DNA templates for sgRNA combining tracrRNA and crRNA were amplified from two complementary ssODNs. sgRNA were transcribed *in vitro* using a MEGA shortscript T7 kit (Thermo Fisher Scientific) and prepared at 2 μg/μl. Linear homology-dependent repair templates (HDRT) containing 35-nucleotide homology arms were amplified using Phusion HF PCR reactions and were prepared at 1 μg/μl. The *dpy-10* co-CRISPR approach^83^, which generates a roller phenotype was used for screening progeny with high edit frequency. A total 15 μl mix of ribonucleoprotein complexes containing Cas9 endonuclease, sgRNA and HDRT for the targeted genes and *dpy-10* were incubated at 37 °C for 10 min and spun at 13,000 rpm for 2 min before gonad microinjections. To generate the sCherry::unc-36 transgenic strain, linear HDRT were amplified using plasmids pcDNA3.1-sfCherry2(1-10)^84^ (Addgene) as PCR templates. For generating 5xMYC::unc-10 and HA::unc-2 strains or mutating the marker gene *dpy-10*, ssODNs (IDT) were used as the HDRT. A silent mutation for the fifth Serine of UNC-10 was introduced in the HDRT to prevent Cas9 recognition. All generated worm lines were outcrossed at least three times to remove potential off-target mutations. A full list of primers, sgRNAs and HDRT is available in Supplementary Table 2.

### Microinjection

Microinjection was performed as previously described^30^. For genome editing, standard gonad microinjection was done on young adult worms. For CALM imaging, synthetic complementary peptide sfCherry2(11)-cysteine (NH_2_-C-acp-linker GSGGGSTSYTIVEQYERAEARHSTGG-COOH, LifeTein, LLC.75% purity) was first dissolved in DMSO to as a 30mM stock, which was then diluted to 750µM with PBS buffer right before microinjections. For immunostaining of extracellular sCherry-UNC-36, rabbit anti-mCherry (Abcam Cat# ab167453, RRID: AB_2571870) was freshly diluted at 1:400 with injection buffer^85^ (20 mM K₂HPO₄, 3 mM potassium citrate, 2% PEG3000, pH 7.4). Either peptide or antibody solutions were spun at 15,000 rpm for 2 min before microinjection into the pseudocoelomic cavity at the head region of staged 1-day adult worms. Animals were recovered in recover buffer (5 mM HEPES pH 7.2, 3 mM CaCl₂, 3 mM MgCl₂, 66 mM NaCl, 2.4 mM KCl, 4% w/v glucose) drop on a NGM plate at room temperature for 16-20 h before imaging or freeze cracking. Microinjection was done with an inverted IX50 Olympus microscope, equipped with an Eppendorf FemtoJet micro-injector connected to FemtotipII microinjection capillaries held on a Leitz manual micromanipulator.

### Behavioral assays

Aldicarb acute paralysis assays and body bending thrashing assays were performed on staged 1-day adult worms as previously described^86,87^. Numbers of body bending per minute were scored in the M9 buffer added to 35 mm sterile NGM agar plates. Worms were first accommodated in M9 buffer for 2 min and the number of sinusoidal curves made within a 2 min period were counted. At least 15 worms were scored for each strain. For aldicarb assays, 6-well NGM plates containing 1 mM aldicarb were made and seeded with OP50 bacteria the day before the assays. For each strain, 30 animals were placed on plates and monitored every 30 min for 4 h. Worms were considered paralyzed if their whole body including nose and tail regions show no movement after a gentle tap on the head using a worm pick. For comparison to mutant animals, wildtype control worms were cultured and scored simultaneously on the same batch of NGM or aldicarb plates. At least three trials, with independent worm cultures, were performed for each assay. Data collections were performed blinded to the animal strains. Data were analyzed using either GraphPad Prism software version 8.0.2(263) or OriginPro software (version 10.1.0.178).

### Image acquisition for CALM

CALM imaging was performed as previously described^30^. 16-20h post peptide microinjection, worms were anesthetized for 6-10 min in M9 buffer containing 30 mg/mL 2,3-butanedione monoxime (BDM, Sigma) in a 35 mm plate. Anesthetized worms were then mounted in 8 μL BDM solution on freshly prepared 2% agarose pads with high-precision Deckglaser microscope coverslips (Marienfeld, 0117650). Worms were positioned with one of the nerve cords facing upright by gentling sliding the coverslip without pressing down. The worm position was secured by sealing the four quarters of the coverslip to the slide with minimal nail polish and stabilized for 10min before imaging. Anesthetized worms were imaged within 40 min, after which locomotion recovered. Simultaneous dual-color imaging of Punc-25::SNB-1::GFP and sCherry-UNC-36 was performed on an inverted Nikon Eclipse Ti-E microscope equipped with HiLO optics, a 100x 1.49 NA objective (Nikon), 488 and 561 nm fiber-coupled excitation lasers (Agilent), a multiband pass ZET405/488/561/647 excitation filter (Chroma), a quad-band ZT405/488/561/647 dichroic mirror (Chroma), two EMCCD cameras (Andor iXon), a dual-camera imaging adapter (Andor), an emission-splitting FF560FDi01 dichroic mirror (Semrock), and emission filters at 525/50 nm (Semrock) for GFP and 600/50 nm (Chroma) for sCherry imaging. Movies were acquired at a frame rate of 80 ms/frame, with EMCCD gains of 300, and 1 x1 binning, leading to image pixel sizes of 107 nm on each camera. Dual-channel alignment was done by imaging 40 nm diameter TransFluoSphere beads (488/685, Invitrogen). CALM imaging of individual sCherry-UNC-36 was performed by continuous HiLO illumination of nerve cords marked by either GFP-UNC-2 or Punc-25::SNB-1::GFP for 1-2 min per field of view from the tail to the head regions of each animal. GFP channel was excited with minimal laser intensity to reduce background. Single molecule detections of sCherry-UNC-36 showed typical blinking events, single-step photobleaching, and sudden fluorescence appearance, as previously described^29,30^. Each strain condition was imaged with at least three independent cultures of worms. The numbers of animals imaged for each strain are indicated in figure legends.

### Tracking and diffusion analyses

Single molecule tracking was performed as previously described^29,30^ using the Matlab software SLIMfast, which is based on multiple-target tracing algorithms^88^. For sub-diffraction limit localizations and trajectory assembly of sCherry-UNC-36, the following parameters were used for all conditions: 107 nm/pixel, 80 ms/frame, 10^−6^ localization error, fixed PSF radius of 1.02 pixel determined by theoretical calculation for a 1.49 NA and 615 nm maximum emission wavelength, 0.5-3 pixels search limit and maximum gap of 3 steps for linking positions. Only trajectories with a minimal number of 3 steps were kept for diffusion analyses. Diffusion analyses of neuronal VGCCs were performed on sCherry-UNC-36 trajectories located along the nerve cords or, in the case of only GABAergic VGCCs, that overlapped with synaptic boutons marked by Punc-25::SNB-1::GFP. For each strain condition, trajectories from multiple worms were pooled together and analyzed using the PDSD method^30^ to differentiate diffusive behaviors, and determine the numbers, fractions and mean square displacements of each diffusion regime. Mean square displacement curves for the first 10 steps were plotted for each behavior and fitted with a 2D circularly confined diffusion model^29^ to compute the diffusion coefficients and radius of confinement. OriginPro software (version 10.1.0.178) was used for data analysis.

### Immunofluorescence staining for confocal and STED microscopy

Worms were harvested at stage 2-day adult or 18h post antibody microinjection for immunostaining of extracellular sCherry-UNC-36. Worms were mounted on slides with ice cold Milli-Q water and frozen on dry ice for 1 hour before freeze-cracking as previously described^89^. Fixation was done overnight at 4 °C in microcentrifuge tubes with phosphate-buffered saline (PBS) containing 1% paraformaldehyde (PFA). For HA-UNC-2 immunostaining, fixation was reduced to 20 min at room temperature. Fixation was followed by three 20 min washes with PBS at 4 °C and blocking in blocking solution (PBS, 5% BSA, 0.5% Triton X-100, 1mM EDTA pH 8, 0.05% Sodium azide) for 1 h at room temperature or overnight at 4 °C. Incubations with primary and secondary antibodies were each done overnight at 4 °C in antibody solution (PBS, 2.5% BSA, 0.5% Triton X-100, 1mM EDTA pH 8, 0.05% Sodium azide). Three 20 min washes with PBST (0.5% Tween) were performed at 4 °C between each incubation and one 5 min final wash with PBS was done before mounting. Samples were mounted using high-precision coverslips (Marienfeld, 0117650) and ProLong Glass Antifade Mountant (Invitrogen). Mounted samples were cured for 24 h at room temperature before imaging. The primary antibodies used were rabbit anti-mCherry (Abcam Cat# ab167453, RRID: AB_2571870; 1:400), mouse anti-MYC (Cell Signaling Technology Cat# 2276, RRID: AB_331783; 1:1000), rat anti-HA (Roche Cat# 11867423001, RRID: AB_390918; 1: 100) and chicken anti-GFP (Aves Labs Cat# GFP-1010, RRID: AB_2307313; 1:200). The secondary antibodies used were goat anti-rabbit STAR RED (Abberior Cat# STRED-1002-500UG, RRID: AB_2833015; 1:400), goat anti-mouse STAR RED (Abberior Cat# STRED-1001-500UG, RRID: AB_3068620; 1:600), goat anti-mouse Alexa Fluor 594 (Jackson Immuno Research Labs Cat# 115-585-146, RRID: AB_2338881; 1:500), goat anti-rat Alexa Fluor 594 (Thermo Fisher Scientific Cat# A-11007, RRID:AB_10561522; 1:200) and goat anti-chicken IgY Alexa Fluor 594 (Thermo Fisher Scientific Cat# A-11042, RRID:AB_2534099; 1:500).

### Image acquisition and analyses for confocal and STED microscopy

Both confocal and STED imaging were performed on a Nikon Eclipse FN1 upright microscope equipped with an Abberior STEDYCON system, four excitation lasers (640, 561, 488, and 405 nm), a pulsed STED laser of 775 nm, and three avalanche photodiode detectors operating in single photon counting mode. 2D STED images were acquired using a 100× Nikon Plan APO 1.45 NA oil immersion objective, fixed pixel size of 15 nm, a dwell time set at 10 µs, 5x line accumulation in photon counting mode, and time gating set at 1 ns with a 6 ns width for all channels. Dual-color STED images were acquired using 775 nm depletion beam with the same depletion power for both STAR RED and Alexa Fluor 594 fluorophores. Confocal scans with 1x line accumulation for all channels were done before STED scans, using 5x line accumulation for the same channels. To overlay STED and confocal images of Punc-25::SNB-1::GFP, 5x line accumulation was also used. Acquisition settings for each channel were identical in all conditions. Imaging field of views with sizes of 10 µm or 20 µm x 3µm were used. 5-15 different animals were imaged for each condition. Only dorsal nerve cords were imaged as cell bodies often interfered with imaging at ventral nerve cords.

For figures display, raw STED images were deconvolved with the Huygens Essential version 24.10 software (Scientific Volume Imaging, The Netherlands, http://svi.nl) using the standard mode and theoretical point spread functions of the STED microscope. For quantitative analysis of UNC-36 and UNC-10, raw images were processed with the Savitzky Golay denoise filter^90^ of the Space Noise Reduce plugin in ImageJ/Fiji, setting the parameter order at 7 and the width at 5. Segmentations and measurements of puncta and nanodomains were done with the Particle Sizer plugin^91^ in ImageJ/Fiji, using the single particle mode for puncta and the watershed mode for irregular structures mode for nanodomain. AZs with planar orientation where both UNC-36 and UNC-10 signals could be clearly resolved were manually selected for measurements. Segmentations parameters were optimized using both WT animals and mutants with severe phenotypes and were kept identical for each protein and all conditions. The number of nanodomains and their nearest neighborhood distance (NND) were quantified by treating each nanodomain as a theoretical point spread function to localize its center position. The images were first processed with the Rolling Ball Background Subtraction algorithm^92^ built in ImageJ/Fiji, using with a 10-pixel rolling ball radius, and center localizations and NND measurements were done in the Picasso software^93^. The same settings were used for all conditions. All measurements were done on 2D STED images taken from staged 2-day adult worms. Data analysis and statistics were performed using OriginPro software version 10.1.0.178.

## Notes

### Competing Interest Statement

The authors have declared no competing interest.

### Summary of Updates

Updated Fig.3, Fig. 4, and Fig. 5. Added Fig. 7. Updated references.

