## Supplementary material for "The nanoscale mobility of calcium channels is driven by readily releasable synaptic vesicles to support precise neurotransmission in live *C. elegans*": Supplental figures and tables

### Supplementary figures and tables

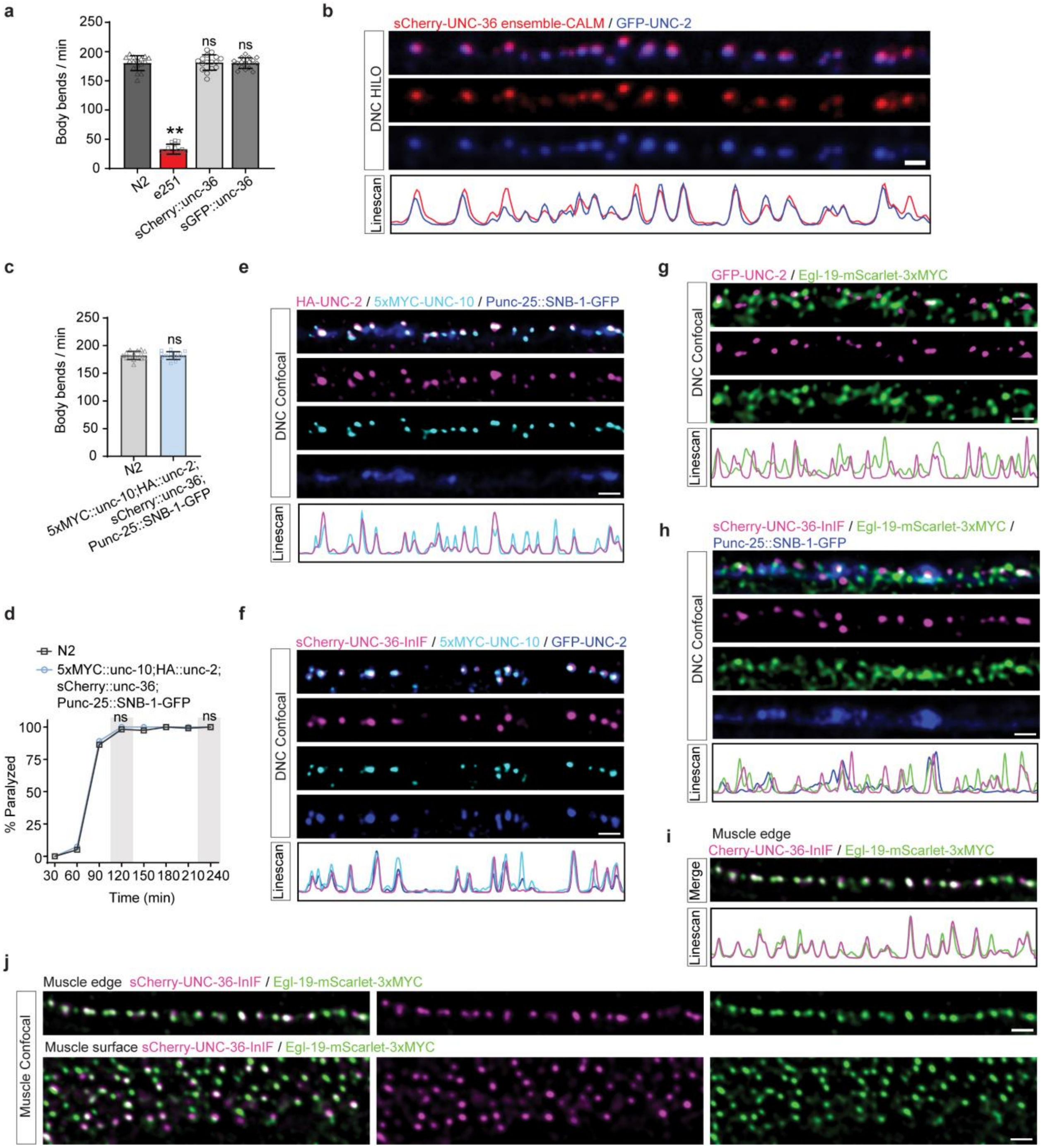

**Figure S1. UNC-36 co-localizes with UNC-2 and UNC-10 at the active zone.**
(a) Trashing assays for animals (n = 16) of indicated genotypes scoring the numbers of body bending per min (mean ± SD). Comparisons with N2 worms by two-tailed t-test, **p<0.01; ns, not significant.

(b) Sum intensity projection of a representative ensemble-CALM movie showing sCherry-UNC-36 (red) co-localized with GFP-UNC-2 (blue) along the dorsal nerve cord (DNC) under HILO illumination. The fluorescence activation of surface sCherry-UNC-36 was maximized in ensemble-CALM by microinjecting excessive amount of sfCherry11 peptides. Linescan through the maxima of sCherry-UNC-36 puncta show the normalized intensity profiles (range: 1-100) of both channels. The Pearson correlation coefficient between sCherry-UNC-36/GFP-UNC-2 is 0.88. Scale bar,1 μm.
(c) Trashing assays for animals (n = 15) of indicated genotypes scoring the numbers of body bending per min (mean ± SD). Comparisons with N2 worms by two-tailed t-test, ns, not significant.
(d) Aldicarb-sensitivity assays measuring the percentage (mean ± SEM) of paralyzed animals over four hours of exposure to 1mM aldicarb. Animals (n = 120) of indicated genotypes were scored together. Each assay was repeated independently four times. Data are mean ± SEM. Comparisons at time point 120 min and 240 min (grey color) by two-way ANOVA and Tukey's tests, ns, not significant.
(e) Representative confocal images of immunostained HA-UNC-2 (magenta) and 5xMYC-UNC-10 (cyan) along a DNC marked by GABAergic Punc-25::SNB-1::GFP (blue). Linescan through the maxima of 5xMYC-UNC-10 shows the normalized intensity profiles (range: 1-100) for each channel. The Pearson correlation coefficient between HA-UNC-2/5xMYC-UNC-10 is 0.77. Scale bar, 1 μm.
(f) Representative confocal images of a DNC showing GFP-UNC-2 (blue) overlayed with immunostained 5xMYC-UNC-10 (cyan) and surface sCherry-UNC-36 (magenta) via antibody microinjection. Linescan through the maxima of sCherry-UNC-36 shows the normalized intensity profiles (range: 1-100) of all three channels. The Pearson correlation coefficient between sCherry-UNC-36/GFP-UNC-2 is 0.83, and between sCherry-UNC-36/5xMYC-UNC-10 is 0.76. Scale bar, 1 μm.
(g) Representative confocal images of a DNC showing GFP-UNC-2 (magenta) and immunostained Egl-19-mScarlet-3xMYC (green). Linescan through the maxima of GFP-UNC-2 shows the normalized intensity profiles (1-100) of both channels. The Pearson correlation coefficient between GFP-UNC-2/Egl-19-mScarlet-3xMYC is 0.20. Scale bar, 1 μm.
(h) Representative confocal images of a DNC marked by Punc-25::SNB-1::GFP (blue) and showing Egl-19-mScarlet-3xMYC (green) and immunostained via anti-MYC and surface sCherry-UNC-36 (magenta) immunostained via anti-mcherry antibody microinjection. Linescan through the maxima of sCherry-UNC-36 shows the normalized intensity profiles (range:1-100) of all three channels. The Pearson correlation coefficient between sCherry-UNC-36/Egl-19-mScarlet-3xMYC is 0.40, sCherry-UNC-36/Punc-25::SNB-1::GFP is 0.24. Scale bar, 1 μm.
(i,j) Representative confocal images of body wall muscles showing a muscle edge and muscle surface immunostained for Egl-19-mScarlet-3xMYC (green) with anti-MYC and for surface sCherry-UNC-36 (magenta) via anti-mcherry antibody microinjection. Linescan through the maxima of sCherry-UNC-36 on the muscle edge in (i) shows the normalized intensity profiles (range: 1-100) for both channels. The Pearson correlation coefficient between sCherry-UNC-36/Egl-19-mScarlet-3xMYC is 0.84. Scale bars, 1 μm.

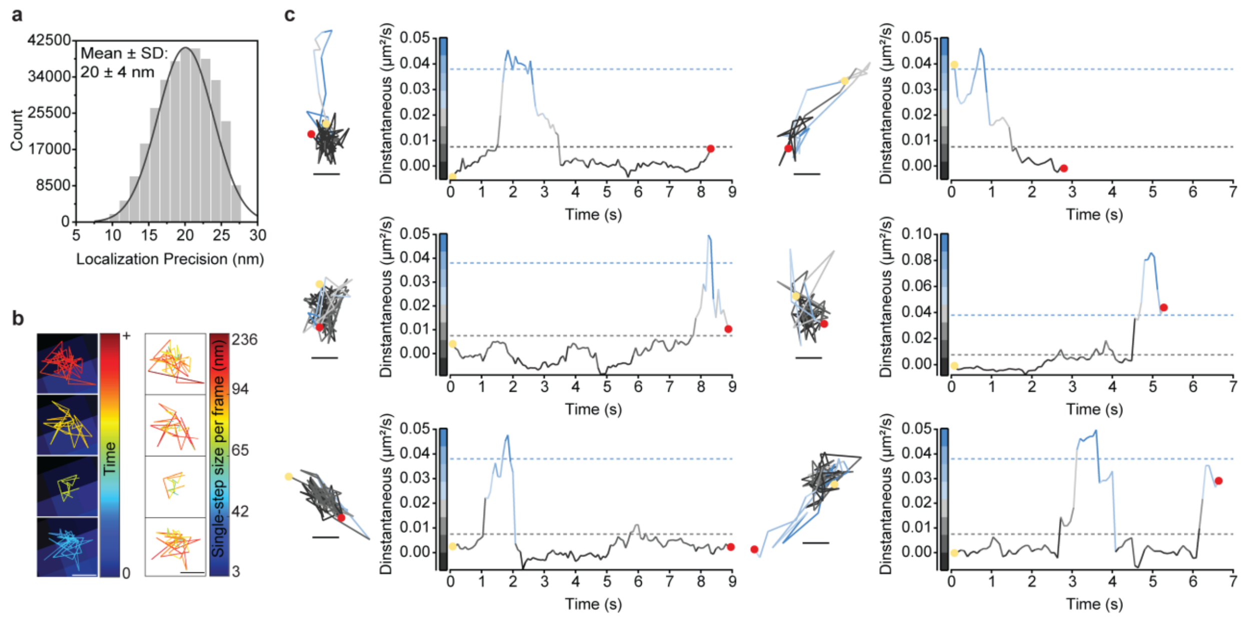

**Figure S2. The two diffusive behaviors of VGCCs identified via PDSD analysis represent transient diffusion states within individual trajectories.**

(a) Distribution of localization precisions for surface sCherry-UNC-36 *in vivo* (n = 322369 emitters; 42,391 trajectories; 58 animals).

(b) Examples of individual VGCC trajectories (figure 1b, asterisk) color coded by time (left; Tmin = blue; Tmax = red) and corresponding color-coding by single-step size per frame (right; min = blue; max = red). Scale bars, 100 nm.

(c) Representative trajectories of heterogeneous VGCC diffusive behaviors color-coded by instantaneous diffusion coefficients (Dinstantaneous). Trajectory start: yellow dot; trajectory stop: red dot; scale bar: 100 nm. Dashed lines reference the $D_{fast}$ (blue) and $D_{slow}$ (black) values extracted from ensemble analysis in figure 1c. Scale bars: 100 nm.

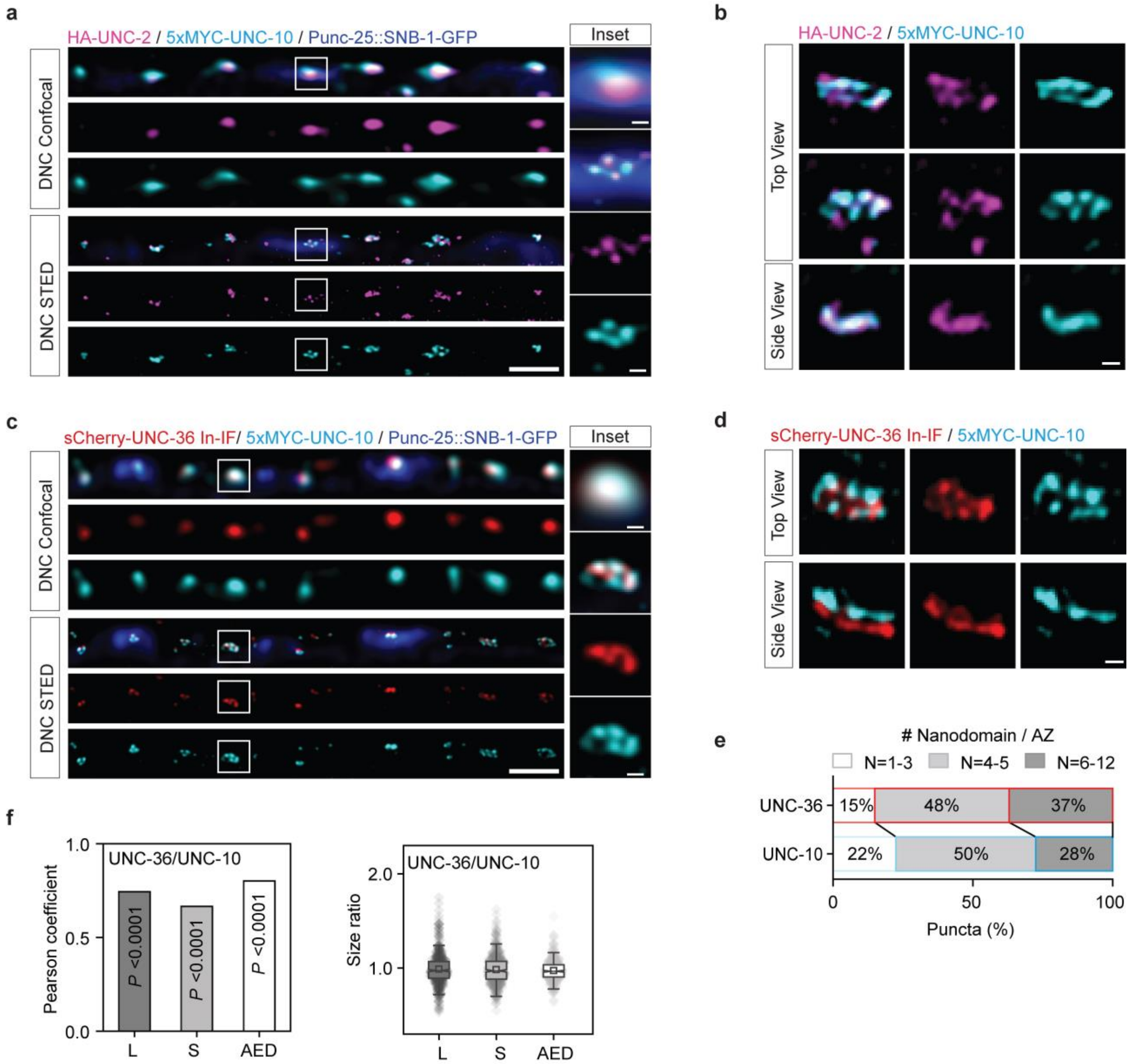

**Figure S3. N-type VGCC complexes and UNC-10 form multiple co-localizing nanodomains within individual AZ puncta**

(a) Representative confocal and dual-color STED images of immunostained HA-UNC-2 (magenta) and
5xMYC-UNC-10 (cyan) along a DNC marked by Punc-25::SNB-1::GFP (blue; confocal only). Scale bars, 1 µm and 100 nm (inset).

(b) Representative top and side views of AZ puncta imaged by dual-color STED and showing HA-UNC-2 (magenta) and 5xMYC-UNC-10 (cyan) nanodomains. Scale bar, 100 nm.

(c) Representative confocal and dual-color STED images of surface sCherry-UNC-36 (red) immunostained by antibody microinjection and of immunostained 5xMYC-UNC-10 (cyan) along a DNC marked by Punc-25::SNB-1::GFP (blue; confocal only). Scale bars, 1 µm and 100 nm (inset).

(d) Representative top and side views of AZ puncta imaged by dual-color STED and showing surface sCherry-UNC-36 (red) and 5xMYC-UNC-10 (cyan) nanodomains. Scale bar, 100 nm.
(e) Percentages of UNC-36 and UNC-10 puncta (n = 516; 10 animals) having different numbers of nanodomains per AZ.
(f) Pearson correlation coefficients (left) and size ratios of the long (L) side length, the short (S) side lengths and the area equivalent circular diameters (AED) between UNC-36 and UNC-10 puncta (n = 516; 10 animals).

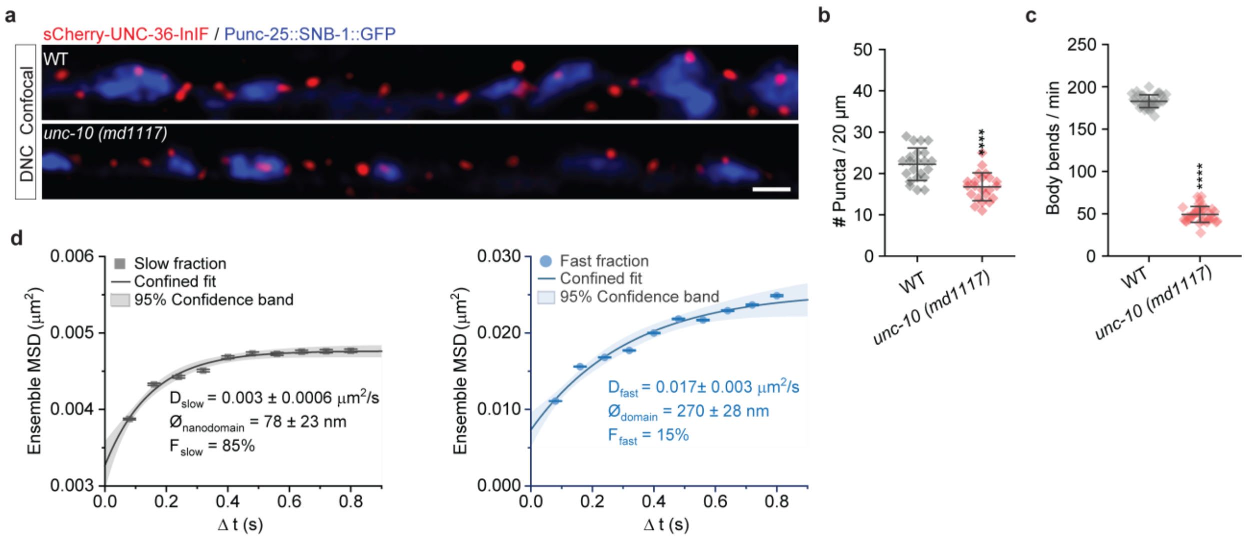

**Figure S4. UNC-10 promotes the synaptic accumulation of VGCCs and enhances VGCC diffusion at AZs.**

(a) Representative confocal images of DNC in WT and *unc-10 (md1117)* mutants showing Punc-25::SNB-1::GFP (blue) and surface sCherry-UNC-36 (red) immunostained via anti-mcherry antibody microinjection. Scale bar, 1 µm.

(b) Number of UNC-36 puncta per 20 µm of DNC (mean ± SD) from WT and *unc-10 (md1117)* mutants (n = 22 images; n = 4 animals). Comparisons by two-tailed t-test, ****p<0.0001.

(c) Trashing assays scoring the numbers of body bending per min (mean ± SD) for WT and *unc-10 (md1117)* mutants (n = 35 animals). Comparisons by two-tailed t-test, ****p<0.0001.

(d) Slow and fast diffusive behaviors of VGCCs extracted from PDSD analyses in *unc-10 (md1117)* mutants (n= 18,645 trajectories; 34 animals). Ensemble MSD ± SEM curves are fitted with a model for 2-dimensional confined diffusion in a circular domain. The 95% confidence bands of each fit are reported, together with the diffusion coefficients (D ± SEM), the diameter of domains (Ø ± localization precision) and the percentage fractions for each slow (black) and fast (blue) diffusive behaviors.

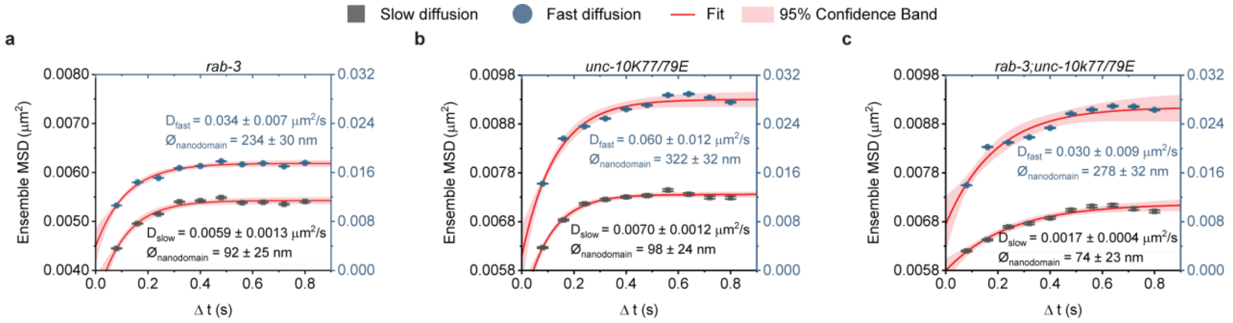

**Figure S5. The mobility and spatial organization of nanodomain-associated VGCCs are regulated by UNC-10/RIM mediated coupling to synaptic vesicles.**

(a-c) Ensemble mean square displacement (MSD ± SEM) curves of slow (black squares) and fast (blue dots) diffusive behaviors extracted from PDSD analyses of VGCCs trajectories in *rab-3(js49)* (a; n = 33,175 trajectories; 107 animals), *unc-10K77/79E (nu487)* (b; n = 29,278 trajectories; 26 animals) and *rab-3(js49); unc-10K77/79E (nu487)* mutants (c; n = 24,455 trajectories; 19 animals). Ensemble MSD ± SEM curves are fitted with a model for 2-dimensional confined diffusion in a circular domain. The 95% confidence bands of each fit are reported, together with the diffusion coefficients (D ± SEM) and the diameter of domains (Ø ± localization precision) for each slow (black) and fast (blue) diffusive behaviors.

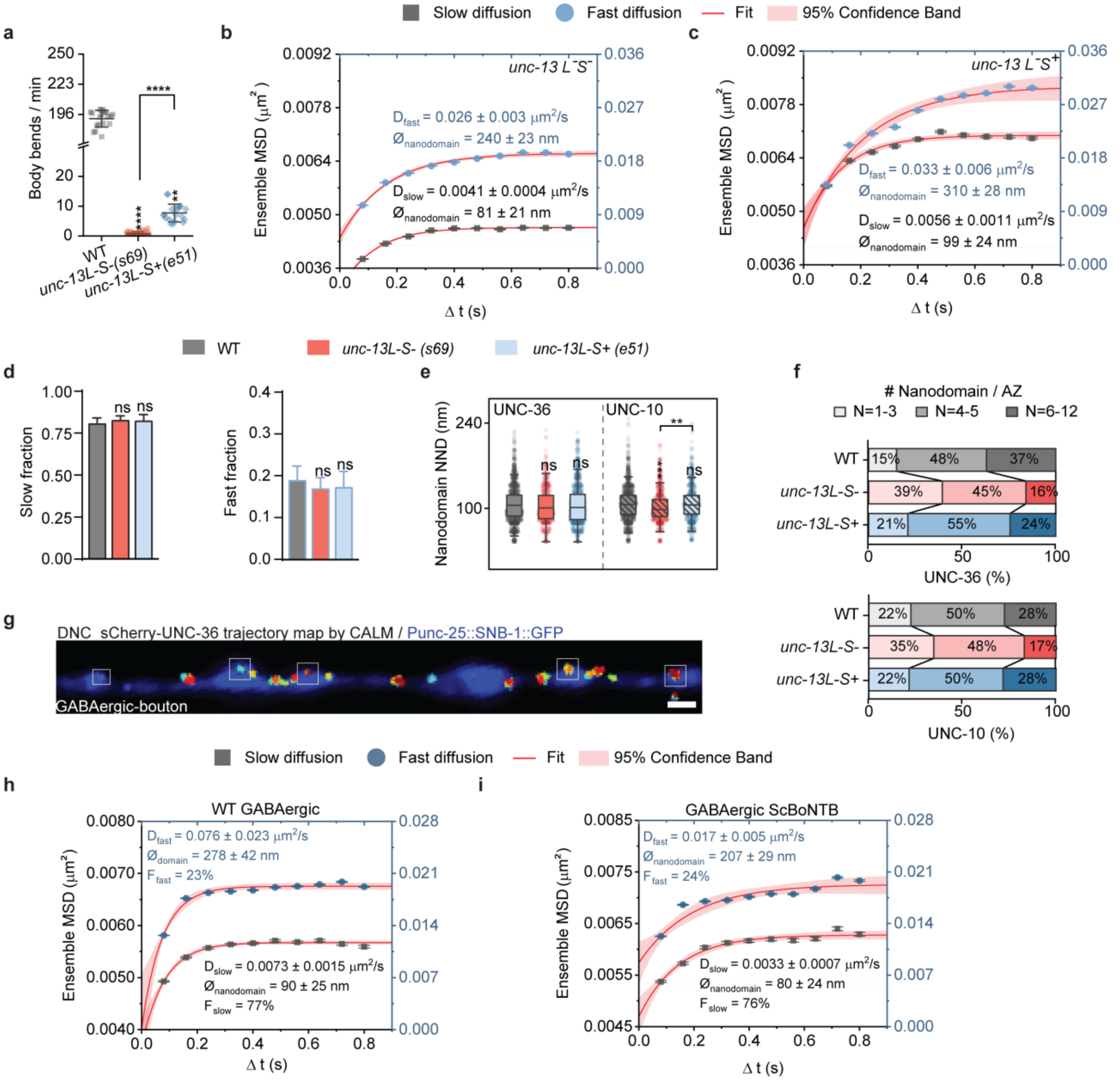

**Figure S6. The mobility and AZ organization of VGCCs is governed by SV priming and SNARE complex assembly.**

(a) Trashing assays scoring the numbers of body bending per min (mean ± SD) for WT (gray; n = 19 animals), *unc-13L-S- (s69)* (red; n = 25 animals) and *unc-13L-S+ (e51)* mutants (blue; n = 19 animals). Comparisons by Kruskal-Wallis and Dunn's test, ****p < 0.0001; **p<0.01.

(b-c) Ensemble mean square displacement (MSD ± SEM) of slow (black squares) and fast (blue dots) diffusive behaviors extracted by PDSD analyses of VGCCs trajectories in *unc-13L-S- (s69)* (b; n = 31,355 trajectories; 35 animals) and *unc-13L-S+ (e51)* mutants (c; n = 19,899 trajectories; 34 animals). Ensemble MSD ± SEM curves are fitted with a model for 2-dimensional confined diffusion in a circular domain. The 95% confidence bands of each fit are reported, together with the diffusion coefficients (D ± SEM) and the diameter of domains (Ø ± localization precision) for each slow (black) and fast (blue) diffusive behaviors.

(d) Fractions (mean ± SEM) for the slow (left) and fast (right) diffusive behaviors extracted by PDSD analyses of VGCCs trajectories from WT (gray; n = 42,391 trajectories; 58 animals), *unc-13L-S- (s69)* (red; n = 31,355 trajectories; 35 animals) and *unc-13L-S+ (e51)* mutants (blue; n = 19,899 trajectories; 34 animals). Comparisons to WT by two-tailed T-test. ns, not significant.

(e, f) Quantifications of nearest-neighbor distance (NND, E) for UNC-36 and UNC-10 nanodomains (e) and percentages of UNC-36 and UNC-10 puncta having different numbers of nanodomains per AZ (f) for WT (gray; n = 516 AZ puncta; 1416 UNC-36 and 1454 UNC-10 nanodomain NND; 10 animals), *unc-13L-S- (s69)* (red; n = 320 AZ puncta; 621 UNC-36 and 718 UNC-10 nanodomain NND; 6 animals) and *unc-13L-S+ (e51)* mutants (blue; n = 205 AZ puncta; 1031 UNC-36 and 816 UNC-10 nanodomain NND; 4 animals). Notched boxes: median ± interquartile range (IQR); whiskers: 1.5xIQR. Comparisons by Kruskal-Wallis and Dunn's test, ***$p < 0.001$; **$p<0.01$; ns, not significant.

(g) Representative image of a DNC with VGCC trajectory maps, outlining the selection criteria for VGCC trajectories located at the tip of GABAergic boutons (white squares). Scale bar, 1 µm.

(h, i) Ensemble mean square displacement (MSD ± SEM) of slow (black squares) and fast (blue dots) diffusive behaviors extracted by PDSD analyses of VGCCs trajectories only at GABAergic boutons in WT animals (h; n = 19,034 trajectories; 42 animals) and ScBoNTB-expressing animals (i; n = 11,549 trajectories; 36 animals). Ensemble MSD ± SEM curves are fitted with a model for 2-dimensional confined diffusion in a circular domain. The 95% confidence bands of each fit are reported, together with the diffusion coefficients (D ± SEM), the diameter of domains (Ø ± localization precision) and the percentage fractions for each slow (black) and fast (blue) diffusive behaviors.

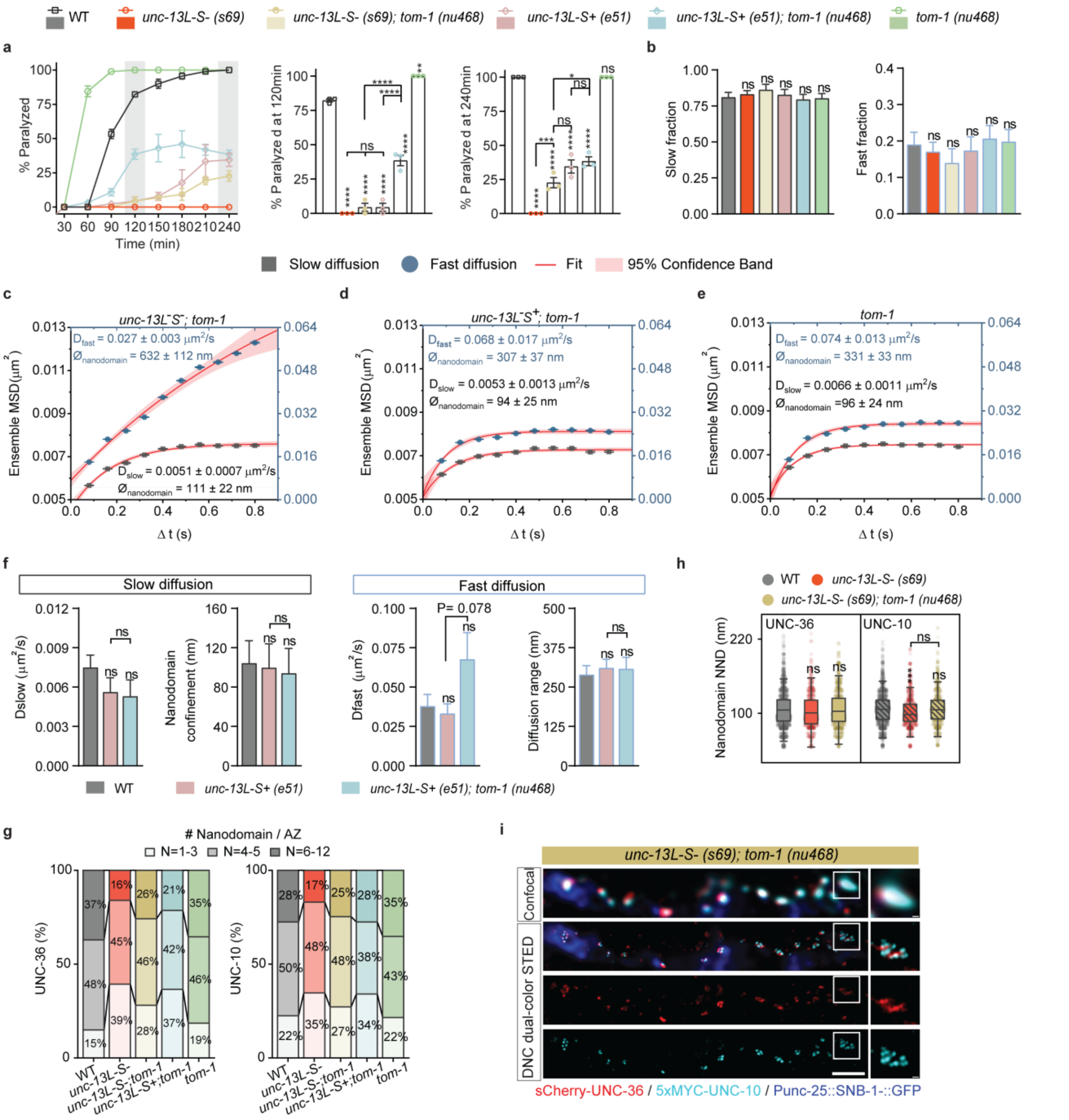

**Figure S7. The distinct mobilities and nanodomain organization of VGCCs are differentially modulated by SV priming levels.**

(a) Aldicarb-sensitivity assays measuring the percentage of paralyzed animals (left; mean ± SEM) over four hours of exposure to 1mM aldicarb. Animals (n = 90; 3 independent tests) of indicated genotypes were scored together. Comparisons to WT at time points or across genotypes at t = 120 min (middle) and t = 240 min (right) by two-way ANOVA and Tukey's tests, ****p<0.0001; ***p < 0.001; **p<0.01; *p<0.05; ns, not significant.

(b) Fractions (mean ± SEM) for slow (left) and fast (right) diffusive behaviors extracted by PDSD analyses of VGCCs trajectories from WT (gray; n = 42,391 trajectories; 58 animals), *unc-13L-S-*

*(s69)* (red; n = 31,355 trajectories; 35 animals), *unc-13L-S- (s69);tom-1(nu468)* (brown; n = 25,309 trajectories; 15 animals), *unc-13L-S+ (e51)* mutants (pink; n = 19,899 trajectories; 34 animals), *unc-13L-S+ (e51);tom-1(nu468)* mutants (cyan; n = 18,963 trajectories; 15 animals), and *tom-1(nu468)* mutants (green; n = 34,062 trajectories; 25 animals). Comparisons with WT by two-tailed T-test. ns, not significant.

(c-e) Ensemble mean square displacement (MSD ± SEM) of slow (black squares) and fast (blue dots) diffusive behaviors extracted by PDSD analyses of VGCCs trajectories in *unc-13L-S-(s69);tom-1(nu468)* (c; n = 25,309 trajectories; 15 animals), *unc-13L-S+ (e51);tom-1(nu468)* mutants (d; n = 18,963 trajectories; 15 animals), and *tom-1(nu468)* mutants (e; n = 34,062 trajectories; 25 animals). Ensemble MSD ± SEM curves are fitted with a model for 2-dimensional confined diffusion in a circular domain. The 95% confidence bands of each fit are reported, together with the diffusion coefficients (D ± SEM) and the diameter of domains (∅ ± localization precision) for each slow (black) and fast (blue) diffusive behaviors.

(f) Mobility of VGCCs in WT (gray; n = 42,391 trajectories; 58 animals), *unc-13L-S+ (e51)* mutants (pink; n = 19,899 trajectories; 34 animals) and *unc-13L-S+ (e51);tom-1(nu468)* mutants (cyan; n = 18,963 trajectories; 15 animals), including diffusion coefficients (mean ± SEM) and nanodomain confinement size or diffusion range (mean diameter ± localization precision) for the channels' slow (left) and fast (right) diffusive behaviors. Comparisons by two-tailed T-test. ns, not significant.

(g) Percentages of UNC-36 (left) and UNC-10 (right) puncta (n = 516; 10 animals) having different numbers of nanodomains per AZ in WT (gray; n = 516 AZ puncta; 10 animals), *unc-13L-S-(s69)* (red; n = 320 AZ puncta; 6 animals), *unc-13L-S- (s69);tom-1(nu468)* (brown; n = 349 AZ puncta; 10 animals), *unc-13L-S+ (e51);tom-1(nu468)* mutants (cyan; n = 276 AZ puncta; 8 animals) and *tom-1(nu468)* mutants (green; n = 529 AZ puncta; 10 animals).

(h) Quantifications of UNC-36 and UNC-10 nanodomain nearest-neighbor distance (NND) in WT (gray; 1416 UNC-36 and 1454 UNC-10 nanodomain NND; 10 animals), *unc-13L-S- (s69)* (red; 621 UNC-36 and 718 UNC-10 nanodomain NND; 6 animals) and *unc-13L-S- (s69);tom-1(nu468)* (brown; 856 UNC-36 and 760 UNC-10 nanodomain NND; 10 animals). Notched boxes: median ± interquartile range (IQR); whiskers: 1.5xIQR. Comparisons by Kruskal-Wallis and Dunn's test, ***$p < 0.001$; ns, not significant.

(i) Representative confocal and dual-color STED images in *unc-13L-S-(s69);tom-1(nu468)* mutants, of surface sCherry-UNC-36 (red) immunostained by antibody microinjection and of immunostained 5xMYC-UNC-10 (cyan), along a DNC marked by Punc-25::SNB-1::GFP (blue, confocal only). Insets show an AZ puncta containing 11 UNC-10 nanodomains. Scale bars, 1 µm and 100 nm (insets).

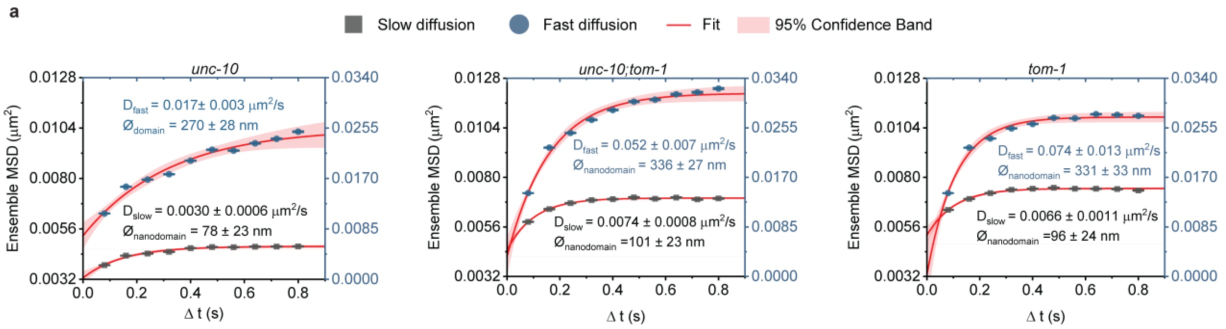

**Figure S8. Primed SVs can directly modulate the mobilities and multi-nanodomain organization of VGCCs downstream of UNC-10.**

(a) Ensemble mean square displacement (MSD ± SEM) of slow (black squares) and fast (blue dots) diffusive behaviors extracted from PDSD analyses of VGCCs trajectories in *unc-10 (md1117)* (left; n= 18,645 trajectories; 34 animals), *unc-10(md1117);tom-1(nu468)* (middle; n = 20,882 trajectories; 26 animals) and *tom-1(nu468)* mutants (right; n = 34,062 trajectories; 25 animals) mutants. Ensemble MSD ± SEM curves are fitted with a model for 2-dimensional confined diffusion in a circular domain. The 95% confidence bands of each fit are reported, together with the diffusion coefficients (D ± SEM) and the diameter of domains (Ø ± localization precision) for each slow (black) and fast (blue) diffusive behaviors.

**Supplementary Table S1**

| Experimental models:<br>*C. elegans*/Strain | Reference | Strain Number | Description |
|---|---|---|---|
| unc-36(yzR11)[sCherry::unc-36] III. | This study | YZR11 | Endogenous sCherry(splitCherry1-10)::unc-36 for surface detection CALM imaging |
| unc-36(yz135)[sGFP::unc-36] III. | This study | YZ5135 | Endogenous sGFP(splitGFP1-10)::unc-36 for surface detection by CALM imaging |
| *unc-36(e251)* III. | Brenner, 1974 | CB251 | *unc-36* loss of function mutant |
| unc-36(yzR11)[sCherry::unc-36] III.; juIs1 [Punc-25::snb-1::GFP] IV.; unc-2(yz7415)[HA::unc-2]X.; unc-10(yz145)[5xMyc:unc-10] X.; | This study | YZ9262 | Endogenous sCherry::unc-36,HA::unc-2,5xMyc::unc-10.Wild type strain for CALM, confocal and STED imaging of N-type complexes with AZ marker. |
| *unc-10(md1117)* X. | Liu et al., 2019 | NM1657 | *unc-10* loss of function mutant |
| unc-2(cim104)[GFP::unc-2] X. | Oh et al., 202 | HKK845 | Endogenous GFP::unc-2 |
| egl-19(utx45)[egl-19(A&B)::mScarlet-I-C1::3xMyc] IV.; unc-2(cim104)[GFP::unc-2] X. | PTK lab | PTK479 | Endogenous GFP::unc-2,egl-19(A&B)::mScarlet-I-C1::3xMyc. Wild type strain for confocal and STED imaging of Cav1 and Cav2. |
| unc-36(yzR11)[sCherry::unc-36] III.; unc-2(cim104)[GFP::unc-2] X.; unc-10(yz145)[5xMyc:unc-10] X. | This study | YZ4512 | Endogenous sCherry::unc-36,GFP::unc-2,5xMyc::unc-10. Wild type strain for CALM, confocal and STED imaging of N-type VGCC complexes with AZ marker. |
| juIs1 [Punc-25::snb-1::GFP+ lin-15(+)] IV. | Hallam & Jin, 1998 | CZ333 | GABAergic synaptic marker. |
| unc-36(yzR11)[sCherry::unc-36] III.; egl-19(utx45) [egl-19(A&B)::mScarlet-I-C1::3xMyc] IV. ; juIs1 [Punc-25::snb-1::GFP] IV. | This study | YZ5317 | Endogenous sCherry::unc-36,egl-19(A&B)::mScarlet-I-C1::3xMyc with GABAergic synaptic marker. Wild type strain for confocal and STED imaging of L-type VGCC complexes. |
| unc-36(yzR11)[sCherry::unc-36] III.; juIs1 [Punc-25::snb-1::GFP] IV. | This study | YZ8509 | Wild type strain for sCherry::UNC-36 tracking by CALM. |
| unc-36(yzR11)[sCherry::unc-36] III.; juIs1 [Punc-25::snb-1::GFP] IV.; *unc-10(md1117)*X. | This study | YZ0927 | *unc-10(md1117)* null mutant for sCherry::UNC-36 tracking by CALM. |
| unc-36(yzR11)[sCherry::unc-36] III.; juIs1 [Punc-25::snb-1::GFP] IV.; unc-2(yz7415)[HA::unc-2]X.; *unc-10(md1117)* X.; | This study | DZ2711 | *unc-10(md1117)* null mutant for confocal and STED. |

| | | | |
|---|---|---|---|
| *rab-3(js49)*II.; unc-36(yzR11)[sCherry::unc-36] III.; juIs1 [Punc-25::snb-1::GFP] IV. | This study | YZ0901 | *rab-3(js49)* null mutant for sCherry::UNC-36 tracking by CALM. |
| *rab-3(js49)*II.; unc-36(yzR11)[sCherry::unc-36] III.; juIs1 [Punc-25::snb-1::GFP] IV.; unc-2(yz7415)[HA::unc-2]X.; unc-10(yz145)[5xMyc:unc-10] X. | This study | DZ0116 | *rab-3(js49)* null mutant for confocal and STED. |
| *unc-10(nu487 K77/79E)*X. | H. Liu et al., 2019 | KP7503 | *unc-10 K77/79E* mutation strain |
| unc-36(yzR11)[sCherry::unc-36] III.; juIs1 [Punc-25::snb-1::GFP] IV.; *unc-10 (nu487 K77/79E)*X. | This study | YZ0903 | *unc-10 K77/79E mutation* strain for sCherry::UNC-36 tracking by CALM. |
| *rab-3(js49)*II.; unc-36(yzR11)[sCherry::unc-36] III.; juIs1 [Punc-25::snb-1::GFP] IV.; *unc-10 (nu487 K77/79E)*X. | This study | YZ0913 | *rab-3(js49),unc-10 K77/79E* double mutant for sCherry::UNC-36 tracking by CALM. |
| *rab-3(js49)*II.; unc-36(yzR11)[sCherry::unc-36] III.; juIs1 [Punc-25::snb-1::GFP] IV.; unc-2(yz7415)[HA::unc-2]X.; *unc-10 (nu487 K77/79E)*X. | This study | YZ1303 | *rab-3(js49),unc-10 K77/79E* double mutant for confocal and STED. |
| *unc-13(s69)* I.; unc-36(yzR11)[sCherry::unc-36] III.; juIs1 [Punc-25::snb-1::GFP] IV.; | This study | YZ0942 | *unc-13(s69)* L-S-null mutant for sCherry::UNC-36 tracking by CALM. |
| *unc-13(s69)* I.; unc-36(yzR11)[sCherry::unc-36] III.; juIs1 [Punc-25::snb-1::GFP] IV.; unc-2(yz7415)[HA::unc-2]X.; unc-10(yz145)[5xMyc:unc-10] X. | This study | YZ4203 | *unc-13(s69)* L-S- null mutant for confocal and STED. |
| *unc-13(e51)* I.; unc-36(yzR11)[sCherry::unc-36] III.; juIs1 [Punc-25::snb-1::GFP] IV. | This study | YZ0920 | *unc-13(e51)* L-S+ mutant for sCherry::UNC-36 tracking by CALM. |
| *unc-13(e51)* I.; unc-36(yzR11)[sCherry::unc-36] III.; juIs1 [Punc-25::snb-1::GFP] IV.; unc-2(yz7415)[HA::unc-2]X.; unc-10(yz145)[5xMyc:unc-10] X. | This study | DZ2013 | *unc-13(e51)* L-S+ mutant strain for confocal and STED. |
| zxEx1152[unc-47p::ScBoNTB; myo-2p::mCherry] | Q. Liu et al., 2019 | ZX2489 | GABAergic ScBoNTB expression strain |

| | | | |
|---|---|---|---|
| unc-36(yzR11)[sCherry::unc-36] III.; juIs1 [Punc-25::snb-1::GFP] IV.; zxEx1152[unc-47p::ScBoNTB; myo-2p::mCherry] | This study | YZ8998 | GABAergic ScBoNTB expression strain for sCherry::UNC-36 tracking by CALM. |
| *tomo-1(nu468)*I.; unc-36(yzR11)[sCherry::unc-36] III.; juIs1 [Punc-25::snb-1::GFP] IV. | This study | YZ0924 | *tomo-1(nu468)* loss of fucntion mutant for sCherry::UNC-36 tracking by CALM. |
| *tomo-1(nu468)*I.; unc-36(yzR11)[sCherry::unc-36] III.; juIs1 [Punc-25::snb-1::GFP] IV.; unc-2(yz7415)[HA::unc-2]X.; unc-10(yz145)[5xMyc:unc-10] X. | This study | DZ2410 | *tomo-1(nu468)* loss of fucntion mutant for confocal and STED. |
| *unc-13(s69)* I. ; *tomo-1(nu468)* I.; unc-36(yzR11)[sCherry::unc-36] III.; juIs1 [Punc-25::snb-1::GFP] IV. | This study | YZ0922 | *unc-13(s69),tomo-1(nu468)* double mutant for sCherry::UNC-36 tracking by CALM. |
| unc-13(s69) I. ; *tomo-1(nu468)* I.; unc-36(yzR11)[sCherry::unc-36] III.; juIs1 [Punc-25::snb-1::GFP] IV.; unc-2(yz7415)[HA::unc-2]X.; unc-10(yz145)[5xMyc:unc-10] X. | This study | DZ2211 | *unc-13(s69),tomo-1(nu468)* double mutant for confocal and STED. |
| *unc-13(e51)* I. ; *tomo-1(nu468)* I.; unc-36(yzR11)[sCherry::unc-36] III.; juIs1 [Punc-25::snb-1::GFP] IV. | This study | YZ0940 | *unc-13(e51),tomo-1(nu468)* double mutant for sCherry::UNC-36 tracking by CALM. |
| *unc-13(e51)* I. ; *tomo-1(nu468)* I.; unc-36(yzR11)[sCherry::unc-36] III.; juIs1 [Punc-25::snb-1::GFP] IV.; unc-2(yz7415)[HA::unc-2]X.; unc-10(yz145)[5xMyc:unc-10] X. | This study | YZ4048 | *unc-13(e51),tomo-1(nu468)* double mutant for confocal and STED. |
| *tomo-1(nu468)*I.; unc-36(yzR11)[sCherry::unc-36] III.; juIs1 [Punc-25::snb-1::GFP] IV.; *unc-10(md1117)* X. | This study | YZ0975 | *unc-10(md1117),tomo-1(nu468)* double mutant for sCherry::UNC-36 tracking by CALM. |
| *tomo-1(nu468)*I.; unc-36(yzR11)[sCherry::unc-36] III.; juIs1 [Punc-25::snb-1::GFP] IV.; unc-2(yz7415)[HA::unc-2]X., *unc-10(md1117)* X. | This study | YZ1008 | *unc-10(md1117),tomo-1(nu468)* double mutant for confocal and STED. |

**Supplementary Table S2**

| Oligo name | Sequence |
|---|---|
| gRNA Common oligo-2 | AAAAGCACCGACTCGGTGCCACTTTTTCAAGTTGATAACGGACTAGC CTTATTTAAACTTGCTATGCTGTTTCCAGCATAGCTCTTAAAC |
| gRNA oligo-1 Dpy-10 | GCAGCTAATACGACTCACTATAGG CTACCATAGGCACCACGAG GTTTAAGAGCTATGCTGG |
| gRNA oligo-1 unc-36 | GCAGCTAATACGACTCACTATAG CAAGCAGTTTTAATAAGgag GTTTAAGAGCTATGCTGG |
| gRNA oligo-1 II unc-36 | GCAGCTAATACGACTCACTATAG CAACAAGCAGTTTTAATAAG GTTTAAGAGCTATGCTGG |
| sgRNA HA::unc-2 | GCAGCTAATACGACTCACTATAGG GCGGCGATTGGTATCATcca GTTTAAGAGCTATGCTG |
| sgRNA Myc::unc-10 | GCAGCTAATACGACTCACTATAGG GAGATGAGTTTCTGCTCCGA GTTTAAGAGCTATGCTG |
| HDR Template for Dpy-10 | cacttgaacttcaatacggcaagatgagaatgactggaaaccgtaccgcatgcggtgcctatggtagcggagcttc acatggcttcagaccaacagcctat |
| HDR Template for UNC-10 3XMyc tagging | gtgcctcattaaatataatcatggacgatccgtcgGAGCAGAAACTCATCTCTGAAGAGGA TCTGGAGCAGAAACTCATCTCTGAAGAGGATCTGGAGCAGAAACTCA TCTCTGAAGAGGATCTGgacgatccgAGTatgatgccggatttatcccatttatctgcagaaga |
| HDR Template for UNC-2 HA tagging | ggctcttaccatcactcgatcgattacagccatggATG TACCCATACGATGTTCCAGATTACGCTATGATACCAATCGCCGCATC GGAAATTATACCATC |
| Primers to generate sGFP::unc-36 HDR Template | |
| HDR Template unc-36 Primer-1 Forward | 5‘ cgccacttatgtctccacaacaagcagttttaataagtg 3’ |
| HDR Template unc-36 Primer-2 Reverse | 5' tgtaaatgtgagacaaatgtgttttatcttacctctccttattaaatccttttcatttgg 3' |
| HDR Template unc-36 Primer-1-1 Forward | 5' cgccacttatgtctccacaacaagcag 3' |
| HDR Template unc-36 Primer-2-1 Reverse | 5' tgtaaatgtgagacaaatgtgttttatcttacctctcc 3' |
| Primers to generate sCherry::unc-36 HDR Template | |
| HDRT mPrimer Forward | 5' cgccacttatgtctccacaacaagcagttttaataagtggccagctagcgtaagttta 3' |
| HDRT mPrimer Reverse | 5' gtgttttatcttacctctccttattaaagtcctcgttgtggctggtgatgtccagc 3' |
| sGFP::unc-36/sCherry::unc-36 genotyping Forward | 5' tccaccaaaaaaatcttaattttcagatctctttttgag 3' |

| | |
|---|---|
| sGFP::unc-36/sCherry::unc-36 genotyping Reverse | 5' tgtaaatgtgagacaaatgtgttttatcttacctctcc 3' |
| Myc::unc-10 genotyping Forward | 5' atacaactcttaaaccgaacaaa 3' |
| Myc::unc-10 genotyping Reverse | 5' tcttctgcagataaatggga 3' |
| HA::unc-2 genotyping Forward | 5' tatgggaaacgacggcga 3' |
| HA::unc-2 genotyping Reverse | 5' ggcgtgcgaatttcaaaaa 3' |
